# The effectiveness of selection in a species affects the direction of amino acid frequency evolution

**DOI:** 10.1101/2023.02.01.526552

**Authors:** Hanon Solomon McShea, Andrew L. Wheeler, Peter Goodman, Catherine Weibel, Sawsan Wehbi, Jennifer E. James, Gavin Huttley, Joanna Masel

## Abstract

Nearly neutral theory predicts that species with higher effective population size (*N*_*e*_) are better at purging slightly deleterious mutations. We compare evolution in high-*N*_*e*_ vs. low-*N*_*e*_ vertebrates to reveal subtle selective preferences among amino acids. We take three complementary approaches. First, we fit non-stationary substitution models using maximum likelihood, comparing the high-*N*_*e*_ clade of rodents and lagomorphs to its low-*N*_*e*_ sister clade of primates and colugos. Second, we compared evolutionary outcomes across a wider range of vertebrates, via correlations between amino acid frequencies and the codon adaptation index of species, a proxy for *N*_*e*_. Third, we dissected which amino acids substitutions occurred in human, chimpanzee, mouse, and rat, as scored by parsimony – this also enabled comparison to a historical paper. All three methods agree on amino acid preference under more effective selection. Preferred amino acids are less costly to synthesize and use GC-rich codons, which are hard to maintain under AT-biased mutation. These factors explain 85% of the variance in amino acid preferences. Within highly exchangeable pairs of amino acids, arginine is strongly preferred over lysine, valine over isoleucine, and aspartate over glutamate, consistent with more effective selection preferring a marginally larger free energy of folding. The first two of these preferences, but not the third, match differences between thermophiles and mesophilic relatives. These results reveal the biophysical consequences of mutation-selection-drift balance, and demonstrate the utility of nearly neutral theory for understanding protein evolution.

**Significance statement:** According to the nearly neutral theory of molecular evolution, selection is less able to distinguish between similar alleles in species with lower population size. We identify which amino acids are subject to such weak preferences – these tend to be less costly to make, to use GC-rich codons easily destroyed by mutation, and to be enriched in thermophiles relative to mesophiles. The latter agrees with theories of marginal protein stability under mutation-selection-drift balance.

## Introduction

Tomoko Ohta (1973) proposed that there is evolutionarily significant variation among species in the ability of selection to purge weakly deleterious mutations. According to Ohta’s nearly neutral theory of evolution, species with small “effective population sizes” (*N*_*e*_) are less able to purge slightly deleterious mutations and will thus retain more of them. By estimating *N*_*e*_ from neutral genetic diversity, Lynch and Conery (2003) used the nearly neutral theory to explain why species with smaller effective population sizes have more bloated genomes. Species with larger effective population sizes also show stronger synonymous codon usage bias due to selection (Akashi, 1996; Galtier et al., 2018; S. Subramanian, 2008; Weibel et al., 2024). Here we ask whether differences among species in amino acid frequencies can similarly be explained by the nearly neutral theory of evolution, and if so, what this tells us about the biophysical basis of intrinsic selective preferences.

We use three complementary approaches to test, for the first time, whether differences among species in the effectiveness of selection can predict amino acid frequencies. First and primarily, we fit non-stationary amino acid substitution models on the basis of maximum likelihood (ML). This is a major departure from assumptions of stationarity built into most models of amino acid substitution (Bui et al., 2021; Dang et al., 2022; Le et al., 2012; Le & Gascuel, 2008; Whelan & Goldman, 2001). To reduce the number of fitted parameters, we hold the exchangeability matrix constant at previously estimated values (Qmammal; Bui et al., 2021), while fitting different equilibrium amino acid frequencies for different parts of the tree (Supplementary Figure 1). We fit all 20 amino acid frequencies, going beyond previous work that allowed variation in only one (Muñoz-Gómez et al., 2022) or two (Groussin et al., 2013) parameters to summarize changing amino acid frequencies. We are able to get all amino acid frequencies because the cogent3 code base (Kaehler et al., 2015; Schranz et al., 2008; Verbyla et al., 2013) enables us to group branches together, avoiding the need to estimate 19 free parameters for every branch. We then compare the fitted equilibrium amino acid frequencies within high-*N*_*e*_ (Phifer-Rixey et al., 2012) Glires branches (including both Rodentia and Lagomorphs) to those within low-*N*_*e*_ (Tenesa et al., 2007) Primatomorpha branches (including Primates plus colugos). A difference in the effectiveness of selection between these two sister clades is well supported (Eőry et al., 2010; Halligan et al., 2010; Keightley et al., 2005), and genome data is high-quality.

Second, we look at a broader species range (all vertebrates), to confirm that results from our substitution model method are driven by *N*_*e*_ rather than by other idiosyncrasies of Glires and/or Primatomorpha biology. We examine whether current proteome-wide amino acid frequencies are correlated with *N*_*e*_. We quantify *N*_*e*_ using a recently developed metric, the Codon Adaptation Index of Species (CAIS; Weibel et al., 2024). CAIS captures the fraction of synonymous sites within the genome for which selection can overcome drift. Because the distribution of selection coefficients is likely to be similar for different vertebrate species, CAIS estimates *N*_*e*_. CAIS uses Kullback-Leibler divergence to quantify the degree to which codon bias departs from that expected from the %GC content of the species. It is calculated with respect to a standardized set of amino acid frequencies and therefore, by construction, also avoids confounding with amino acid frequencies. Weibel et al. (2024) found that high-CAIS species have higher proteome-wide intrinsic structural disorder, demonstrating the metric’s utility for revealing how the effectiveness of selection shapes fundamental physical properties of macromolecules. Conveniently, CAIS requires only a single complete genome for its estimation, unlike traditional *N*_*e*_ estimates which require data on polymorphisms and mutation rate. CAIS directly measures the property of interest, namely the degree to which weak selection is able to exert subtle selective preferences within a species.

Third, noting that our ML method only captures net amino acid frequency change, we use parsimony to count instances of change, to determine which amino acid pairs obey detailed balance vs. contribute to overall amino acid frequency change. This parsimony approach is also partially motivated by resolving a historical controversy, after Jordan et al. (2005) used a similar approach to claim universal patterns of amino acid gains and losses across the tree of life, dating back to the origins of the genetic code.

The findings of Jordan et al. (2005) were subject to intense criticism on three counts. First, the method assumed that all derived alleles were substitutions, when some instead could be polymorphisms – many of them slightly deleterious – segregating in the population (Hurst et al., 2006; McDonald, 2006). Our analysis filters out polymorphic sites to obtain a dataset of true substitutions (Figure 1A), something it was not possible to do at the time Jordan et al. (2005) was published. Removing contamination by slightly deleterious polymorphisms should better reveal any true universal trends. To ensure the availability of polymorphism data, we restrict our analysis to two pairs of sister species. We compare recent fluxes in the primates *Homo sapiens* and *Pan troglodytes* to a shared set of sequences in the rodents *Rattus norvegicus* and *Mus musculus*.

**Figure 1:**
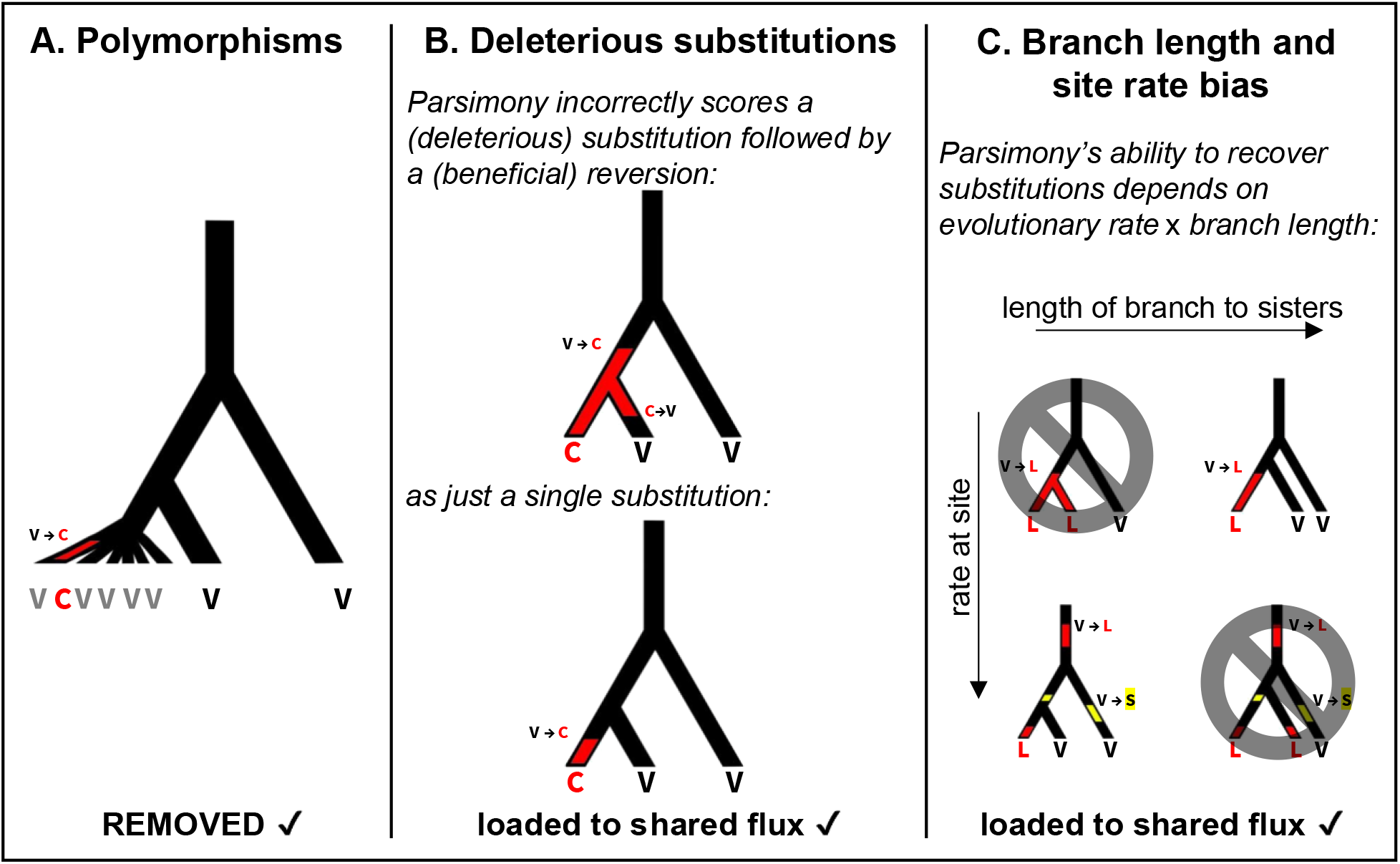
Artifacts of the parsimony net flux method and our approaches to them. Every color change along a branch represents a substitution at a focal site. (A) We removed polymorphisms. (B and C) Deleterious substitutions and branch length issues are expected to affect rodent and primate branches similarly. We therefore subtracted flux that was seen in both rodent and primate branches, leaving a metric that describes how rodent flux differs from primate flux (Figure 3C). V→C is used as an example of a slightly deleterious mutation prone to reversion, while V→L and V→S are used as examples of effectively neutral substitutions. On the left side of (C), when sister branch lengths are short, there will be greater sampling of fast-evolving sites, and when sister branch lengths are long, it is easier to detect slow-evolving sites, because they are less prone to multiple events, both within the long sister branches and in the still longer outgroup branch

A second potential problem with this parsimony-based method is that even a set of true substitutions will be enriched for slightly deleterious substitutions (McDonald, 2006). This is because the brevity of their fixed status makes them more likely to survive ascertainment bias when in the derived state than when in the ancestral (Figure 1B). According to this argument, removing slightly deleterious polymorphisms should retain the direction of flux while reducing its magnitude; slightly deleterious polymorphisms should mirror slightly deleterious fixations. A similar argument applies with respect to equilibrium amino acid frequencies; a rare amino acid is comparable to a deleterious allele (McDonald, 2006), explaining earlier, similar results by Zuckerkandl et al. (1971), who observed an apparent increase in rare amino acids.

Goldstein and Pollock (2006) noted a third artifact that depends on the pattern of branch lengths, combined with rate heterogeneity (Figure 1C). In their simulations, the length of the branch between sister species relative to their branch with the outgroup affected which types of substitutions were parsimony-informative and thus detectable. This bias in detection can produce the appearance of strong net flux, even in simulations of a stationary and time-reversible model of protein evolution, for which net flux is by definition zero. Because Jordan et al. (2005) selected their triplets of species on the basis of similar branch length patterns (sister taxa pairwise amino acid divergence <15%, outgroup <25%), as do we, the same branch-length artifact will appear within any trends shared across different species triplets. Our approach is to statistically differentiate taxonomically specific patterns in flux from “shared flux” values that capture both this artifact and that from slightly deleterious fixations, as well as any true universal trends.

Here we fit non-stationary models of amino acid evolution to discover which amino acids are favored by weak selection that is effective only in higher *N*_*e*_ species. We compare to parsimony-based results to resolve a historic controversy, and to obtain a detailed picture of which amino acid substitutions contribute to the non-stationarity. We confirm these flux results across a broader taxonomic range by describing how amino acid frequency outcomes correlate with *N*_*e*_.

## Results

### Low *N*_*e*_ results in costly amino acids produced by mutation bias

Cheaper amino acids are at higher equilibrium frequencies in the higher *N*_*e*_ clade, according to our maximum likelihood method (Figure 2A, Pearson’s R^2^ = 0.76, p = 7x10^-7^). GC-rich amino acids are also retained to a greater degree under more effective selection (Pearson’s R^2^ = 0.40, p = 0.003). The latter is consistent with more effective selection being necessary to overcome the pervasive AT bias of mutation (Haag-Liautard et al., 2008; Hershberg & Petrov, 2010; Lynch, 2010; Lynch et al., 2008), including but not limited to methyl-cytosine deamination, which makes the retention of GC-rich amino acids difficult (although GC-biased gene conversion might partially counter this). GC-rich amino acids are also favored in highly expressed (R^2^ = 0.27, p = 0.02; Jansen & Gerstein, 2000) and slowly evolving (R^2^ = 0.20, p = 0.05; Cherry, 2010) proteins. However, across all 20 amino acids, we do not find a significant correlation between our ML equilibrium difference scores and these papers’ estimates of amino acid enrichment as a function of expression or evolutionary rate.

**Figure 2:**
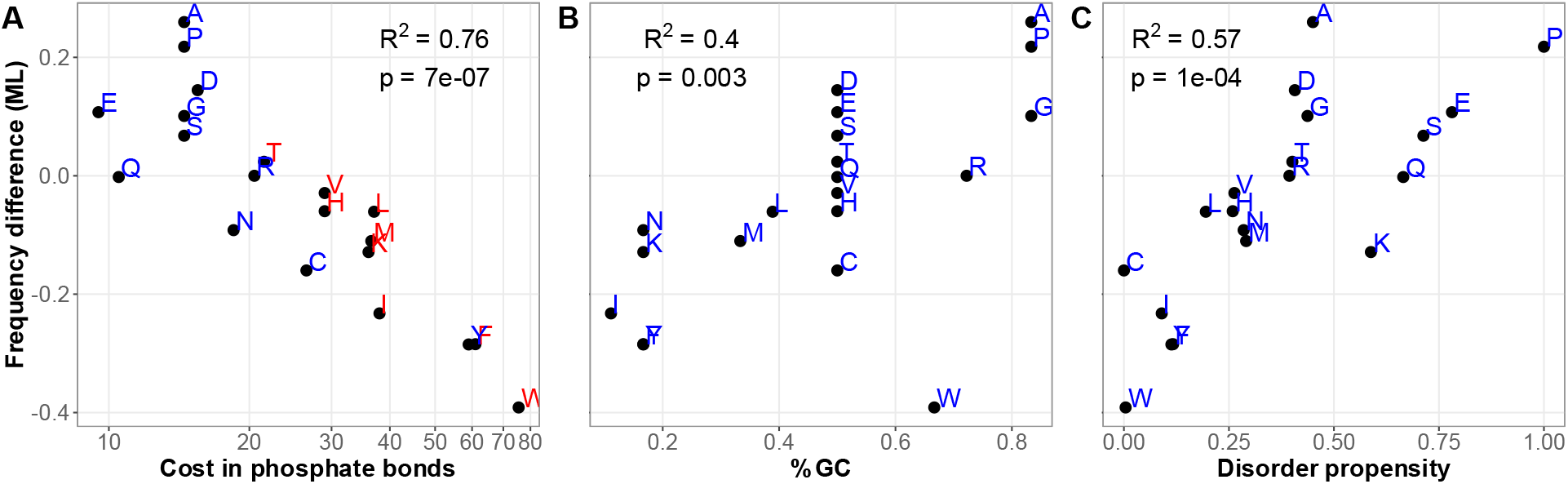
The amino acids preferred in Glires vs. Primatomorpha evolution are cheaper, higher %GC, and more prone to structural disorder. The y-axis “Frequency difference (ML)” indicates equilibrium logit(Glires frequencies) – logit(Primatomorpha frequencies) estimated under a non-stationary model of amino acid substitution (numbers in Supplementary Table 1). This model was fitted by maximum likelihood to an aligned set of orthologs, with one set of equilibrium amino acid frequencies fitted to branches within a clade of 12 Glires species, another to branches within a sister clade of 16 Primatomorpha species, and a third to their shared root (see Methods). The time-reversible exchangeability matrix (Q.mammal; Bui et al., 2021) was held constant. Error bars show ± 1 standard error; R and p-values on figure panels are for Pearson’s R correlation. (A) Biosynthetic cost in *Saccharomyces cerevisiae* under aerobic conditions, as a count of high-energy phosphate bonds (Raiford et al., 2008); we obtain similar R^2^ = 0.77 and p = 4x10^-7^ for aerobic cost on a linear scale. Colors show essentiality in animals (Costa et al., 2015). (B) Disorder propensity (Theillet et al., 2013). (C) %GC, calculated as average %GC for each amino acid’s set of codons in the standard genetic code.

We expect a stronger mutation bias toward AT (and/or weaker counteracting GC-biased gene conversion in low-recombination genomic regions) to create a stronger relationship between an amino acid having GC-rich codons and being used more by higher *N*_*e*_ species. Indeed, as shown in Supplementary Figure 2, the relationship is stronger when we analyze only the third of genes with the lowest %GC at synonymous sites in humans (0.21 < synonymous GC < 0.43, Spearman’s R^2^ = 0.47), than in the middle quantile (0.43 < synonymous GC < 0.54, R^2^ = 0.44) or high-GC quantile (0.54 < synonymous GC < 0.90, R^2^ = 0.31). It is notable that the direction of the relationship does not flip, but only weakens, when the analysis is restricted to high-GC genes. This is compatible either with high-GC amino acids being intrinsically superior, or as discussed in Weibel et al. (2024), that local GC content is less informative than global GC content, because local GC content changes rapidly relative to the amino acid biases studied.

Other predictors of equilibrium frequency differences are significant on their own but drop out of multiple regression models. Most notably, the degree to which an amino acid promotes structural disorder is predictive on its own (Figure 2C, Pearson’s R^2^ = 0.57, p = 0.0001), but drops out of a multiple regression (R^2^ = drops from 0.88 to 0.85 when disorder is removed; p = 0.09 in multiple regression, disorder slope falls from 0.48 to 0.14 in single vs. multiple regression) that retains cost (p = 0.0002, slope drops in magnitude from -0.008 to -0.006) and %GC (p = 0.009, slope falls from 0.47 to 0.21). Disorder is confounded with cost (Pearson’s R^2^ = 0.48, p = 0.0007), making causation hard to pin down.

Similarly, amino acids preferred in the higher *N*_*e*_ clade also tend to have smaller volume (Pearson’s R^2^ = 0.57, p = 0.0001), to be dietarily non-essential for animals (using Costa et al., 2015’s conservative list of essential amino acids) (Figure 2A; t-test p = 0.02), to have lower “stickiness,” an empirical measure of an amino acid’s tendency to be in protein-protein interfaces rather than in non-interface surfaces (Pearson’s R^2^ = 0.36, p = 0.005; Levy et al., 2012), and to have higher polarity (Pearson’s R^2^ = 0.29, p = 0.01). They have more positive marginal fitness effects when expressed as part of a random peptide in the *Escherichia coli* experiments of Neme et al. (2017), as inferred by Kosinski et al. (2022) (Pearson’s R^2^ = 0.39, p = 0.003), as expected given the dependence of these marginal fitness effects on size and disorder, both predictive themselves. These variables, like structural disorder, are confounded with cost and drop out of multiple regressions that also have cost as an independent variable. While selection on volume, essentiality, stickiness, polarity, and structural disorder cannot be completely excluded, the strongest statistical support is for weak selection against more expensive amino acids.

### Parsimony fluxes shared by rodents and primates are likely driven by artifact

According to our parsimony method, mice and rats have very similar net amino acid fluxes to each other (Supplementary Figure 3), as do humans and chimpanzees. This is expected, since %GC content, effective population size, and many other possible influences are similar within each closely related species pair. We therefore calculated “rodent” flux values by combining the amino acid gain and loss counts for mouse and rat branches, and “primate” flux values by combining the counts for human and chimpanzee branches (numbers in Supplementary Table 2).

While our rodent and primate net fluxes are similar to those of Jordan et al. (2005), they are generally of smaller magnitude, particularly in primates (Figures 3A and 3B). We interpret this as a weaker version of the same bias toward slightly deleterious mutations within our fluxes compared to those of Jordan et al. (2005). As discussed in the Introduction (Figure 1B), fixation events scored by parsimony are also expected to be biased toward slightly deleterious mutations. This is because slightly deleterious mutations might sometimes reach fixation but will revert relatively quickly to a less deleterious state along subsequent branches, and therefore will be scored incorrectly.

**Figure 3:**
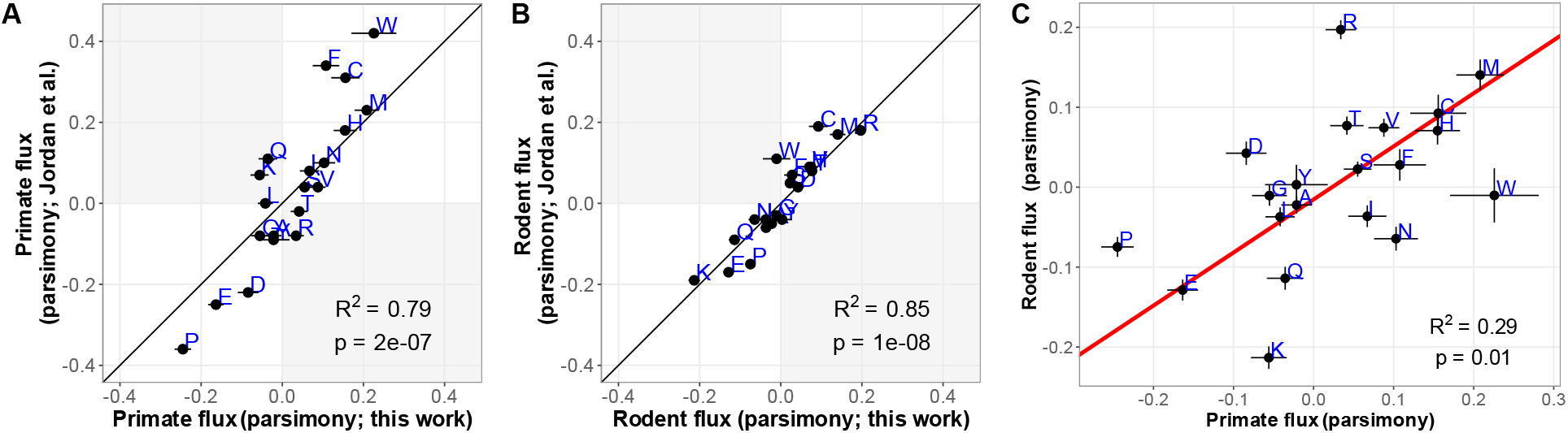
Removing polymorphisms reduces the magnitude of parsimony flux, especially for primates (A and B). Net fluxes depend less on taxon when calculated by parsimony (C) than by maximum likelihood (D). The parsimony-based net flux for each amino acid = (# gains – # losses) / (# gains + # losses), making values greater than 0 indicate a tendency towards gain. Error bars show ± 1 standard error; R^2^ and p-values on figure panels are for Pearson’s R correlation. (A and B) Black diagonal lines show y = x, shaded areas show sign reversal. (C) Red line shows the fit of a type II linear regression (y = 0.664x – 0.0154). Shared flux and flux difference are calculated from this regression line, as discussed in the main text.

The main difference between our method and that of Jordan et al. (2005) is that we exclude polymorphisms. Our primate species pair is more closely related than our rodent species pair, i.e. it has fewer true substitutions. If we assume similar levels of polymorphisms in both, the smaller true divergence in primates means that removing polymorphism has a proportionately larger impact on primate fluxes than rodent fluxes. This is compatible with the observed greater deflation in net flux magnitudes that we see for primates (Figure 3A) than for rodents (Figure 3B).

Remaining artifacts in our polymorphism-free version of the parsimony method should be shared between rodents and primates (in addition to true universal trends, if any). In contrast, differences in net parsimony fluxes may reflect the more effective selection in rodents relative to the mutation bias that overwhelms selection in primates. By statistically isolating shared vs. species-specific patterns in flux, any universal trend or artifact should be reflected in the shared component, while the species-specific component will allow us to estimate the degree to which each of the 20 amino acids is favored vs. disfavored in more effectively adapted rodents relative to less well adapted primates. To do this, we fit a type-II linear regression model (Figure 3C, red line) and calculated the projection along the model line (“shared flux”) for each amino acid, as well as the shortest distance from that line (“flux difference”). The shared component thus reflects fluxes that are common to both rodents and primates, and the flux difference reflects subtle differences between the patterns of fluxes in the two groups. The sign we give to the flux difference allows it to be interpreted as a rodent-driven component, i.e., a positive value indicates that the amino acid is gained in rodents to a greater degree (or lost to a lesser degree) than in primates.

As expected for a successful partition, the shared component of parsimony fluxes has a strong positive relationship with Jordan et al.’s (2005) average flux (Figure 4A), while the flux difference component has none (Pearson’s R^2^ = 0.03, p = 0.4). Results are similar albeit slightly weaker for Jordan et al.’s (2005) more complex gain/loss rate metric. Shared flux captures the gain of rarer amino acids (Figure 4B) expected from regression to the mean, as observed by Zuckerkandl et al. (1971).

**Figure 4:**
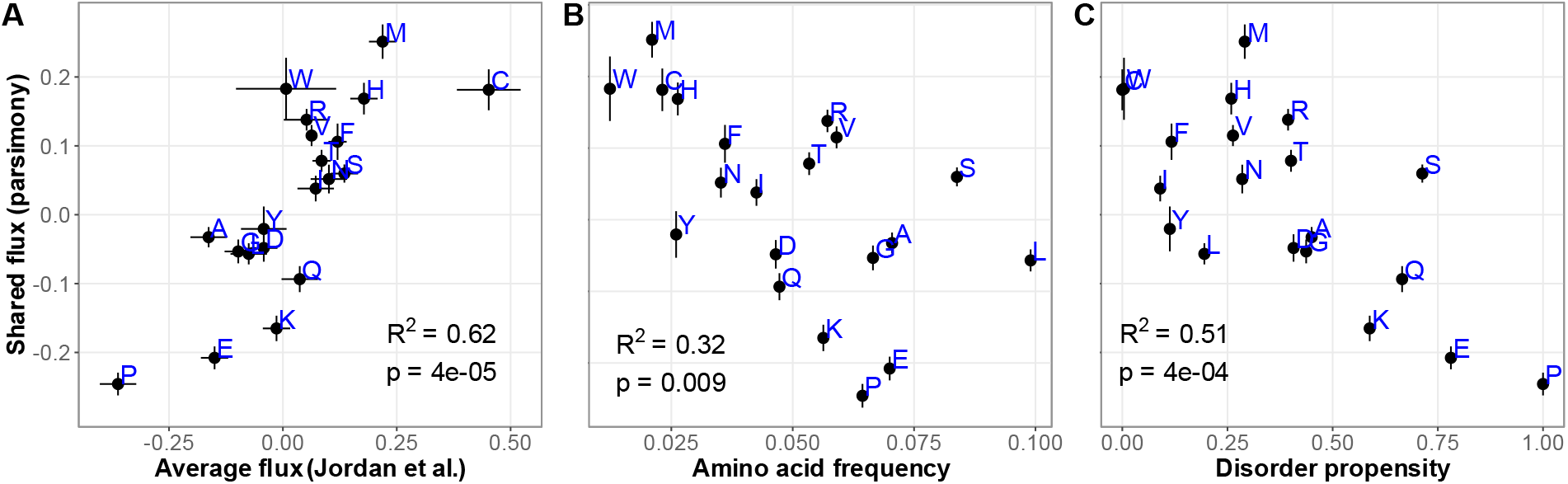
The slightly deleterious mutations artifactually captured by the parsimony method tend to reduce intrinsic structural disorder. (A) Jordan et al. (2005) capture something similar to our shared flux metric. (B) Shared flux describes an increasing frequency of rare amino acids. Frequency is measured as the average of human and chimpanzee amino acid frequencies, with similar results (Pearson’s R^2^ = 0.30, p = 0.01) for the average of mouse and rat amino acid frequencies. (C) Shared flux describes substitutions that reduce structural disorder. Disorder propensities of amino acids were measured by Theillet et al. (2013) on the basis of overrepresentation in disordered vs. well-folded proteins and protein regions. Error bars show ± 1 standard error; R^2^ and p-values on figure panels are for Pearson’s correlation.

Jordan et al. (2005) claimed that the universal trend represented the slow gain of amino acids added late to the genetic code. We find the same relationship between shared flux and the hypothesized order of amino acid recruitment into the genetic code from Trifonov (2000; Pearson’s R^2^ = 0.24, p = 0.03) that Jordan et al. found for their universal trend. However, when we used the revised order of amino acid recruitment inferred by Wehbi et al.’s (2024) “LUCA usage”, we find no correlation with the results of Jordan et al. (2005), nor with our shared flux (p = 0.7 for both, weights based on standard errors on LUCA usage). This is unsurprising given that not all protein-coding sequences can be traced back through recognizable sequence homology to the early days of the genetic code (James et al., 2021; Van Oss & Carvunis, 2019; Wehbi et al., 2024).

If the shared flux reflects the slightly deleterious substitutions artifact of Figure 1B, we expect it to correlate with markers of being slightly deleterious from Figure 2, in particular cost and %GC. However, shared flux does not predict cost or %GC. We note that a correlation between a marker of being slightly deleterious and primate-specific flux could cause an artifact from Jordan et al. to fail to load onto shared flux.

Shared flux does describe decreasing disorder (Figure 4C; Pearson’s R^2^ = 0.62, p = 4x10^-5^), and the related measures of increasing stickiness (Pearson’s R^2^ = 0.40, p = 0.003), increasing mean relative solvent accessibility (RSA) (Pearson’s R^2^ = 0.37, p = 0.004; Tien et al., 2013), and increasing polarity (R^2^ = 0.31, p = 0.01). This is interesting in light of the more subtle branch length ascertainment bias illustrated in Figure 1C. If the shared component of parsimony flux preferentially represents fast-evolving sites (Figure 1C), we note that these are enriched in disordered regions, where slightly deleterious mutations will tend to decrease disorder through regression to the mean. The fact that Jordan et al.’s average flux corresponds to falling disorder is thus consistent with the artifact of Figure 1C, or with synergy between the two artifacts of Figures 1B and 1C.

### Three methods agree on which amino acids are preferred by selection

The slopes of amino acid frequencies as a function of effective population size (Supplementary Table 3) describe outcomes rather than fluxes, across a larger range of vertebrate species. We use the Codon Adaptation Index of Species (CAIS) as a proxy for effective population size, capturing the proportion of codons subject to effective selection, while controlling for GC and amino acid frequencies (see Methods). Supplementing the among-species evidence for the species-wide CAIS measure given by Weibel et al. (2024), we show in Figure 5 that CAIS, when measured for genes rather than species, is correlated with the expression level of highly expressed genes (as is the more commonly used Codon Adaptation index (CAI; Sharp et al., 2010; Sharp & Li, 1987)). In both mouse and humans, the correlation applies only to the 15% most highly expressed genes – this cutoff was chosen from inspection of a smoothed curve (Figure 5). However, both the correlation coefficient and the slope are larger for mouse than humans, reflecting selection at a larger fraction of sites in mouse within these highly expressed 15% genes. This validates that CAIS captures differences between species in the proportion of synonymous sites under effective selection. This also helps validate prior use of CAI just on highly expressed genes (Sharp et al., 2005; Subramanian, 2008), and confirms that mammals are subject to selection on codon usage, adding to the evidence presented by Yang & Nielsen (2008; Table 3), Subramanian (2008; Figure 1C), and Bénitière et al. (2025; Figure 4D).

**Figure 5:**
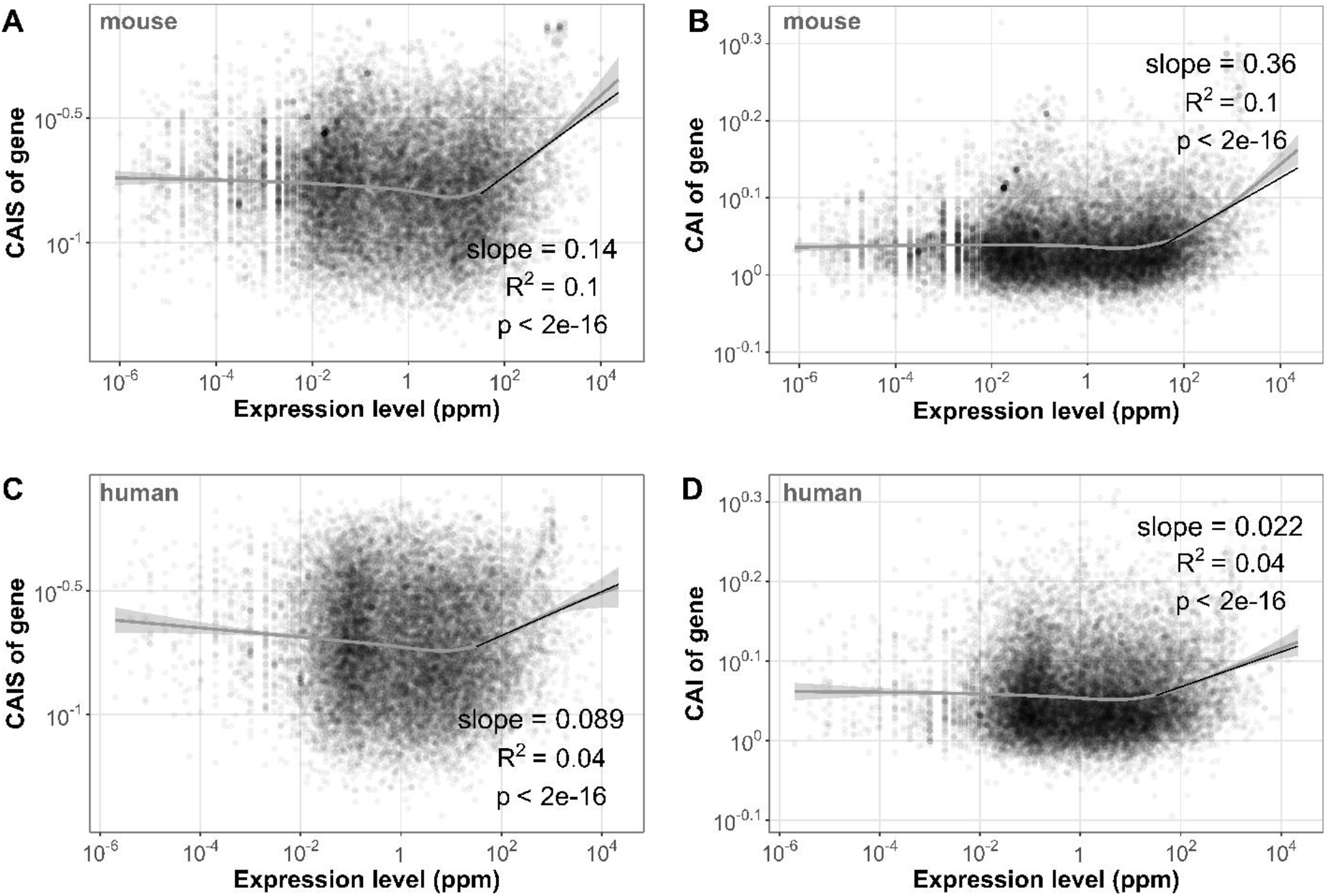
CAIS and CAI depend more strongly on expression level in mouse than in human. Generalized additive models (gray line; implemented in R function geom_smooth and shown with 95% confidence interval) were fit in order to visually determine a threshold for high expression (10^1.5^), above which a linear model (black line) was fit. Pearson’s R^2^ and p values are shown for the linear model.

Our three different metrics of selective preferences among amino acids are in good agreement (Figure 6). They also reveal marked selective preferences between biophysically similar and highly exchangeable amino acids (pairs joined by arrows in Figure 6, discussed in detail below). Our CAIS-based vertebrate preference measure and flux measure will both tend, by construction, to amplify selective preferences between similar amino acids in a way that our ML method will not, accounting for the steepness of the slopes of these lines in Figures 6B and 6C; this reduces Pearson’s R^2^.

**Figure 6:**
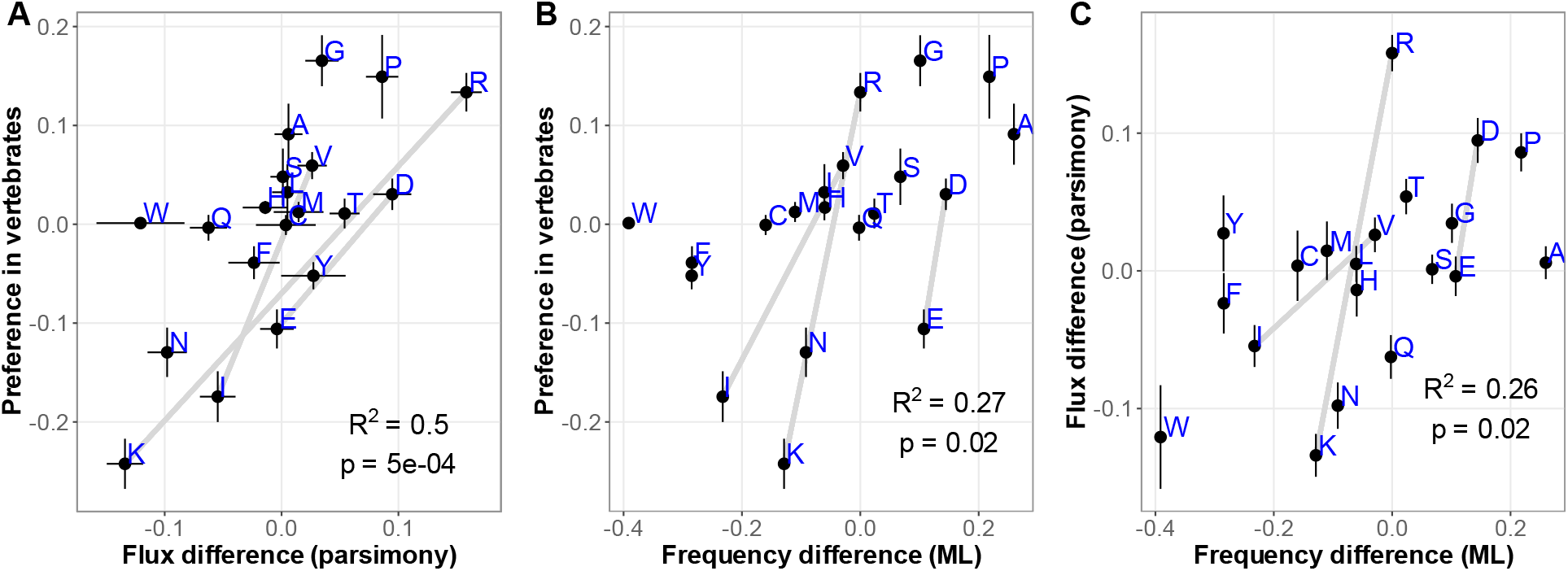
Three distinct methods with different biases recover similar selective preferences among amino acids. “Preference in vertebrates” indicates slopes from linear regressions of amino acid frequency on CAIS. “Flux difference (parsimony)” indicates the shortest distance of an amino acid to the model II fit shown in Figure 2. “Frequency difference (ML)” indicates logit transformed equilibrium Glires frequencies – logit transformed equilibrium Primatomorpha frequencies estimated under a non-stationary model of amino acid substitution. Error bars show ± 1 standard error; R^2^ and p-values are for Pearson’s correlation. Connected pairs are biophysically similar, highly exchangeable amino acids for which all metrics show a marked preference in the same direction.

As is the case for our main metric of selective preference (ML equilibrium frequency difference; discussed above), codon %GC predicts parsimony flux difference (Pearson’s R^2^ = 0.23, p = 0.03) and preference in vertebrates (Pearson’s R^2^ = 0.69, p = 5x10^-6^). Preference in vertebrates is also predicted by mean entropic penalty of folding (Pearson’s R^2^ = 0.23, p = 0.03; Doig & Sternberg, 1995) and marginal fitness in *E. coli* (Pearson’s R^2^ = 0.38, p = 0.004). However, the latter is only explained by Kosinski’s model residuals (Pearson’s R^2^ = 0.22, p = 0.04) – unlike ML frequency difference, neither preference in vertebrates nor parsimony flux difference is significantly predicted by cost or %GC. These results are consistent with our ML measure being superior to the other two, but given broad agreement, we nevertheless use the other two metrics to conduct follow-up analyses not easily possible via ML.

### Different kinds of protein domains have mostly similar amino acid preferences

Applying our CAIS method separately to subsets of the data shows that vertebrate amino acid preferences are similar in ancient vs. recently born Pfam (Mistry et al., 2021) domains (Figure 7A). Partial exceptions are preferences against lysine and glutamic acid, and preferences for leucine and glycine, both of which are stronger in young than in old Pfams. Transmembrane vs. non-transmembrane domains also have similar amino acid preferences, despite operating in different biophysical contexts (Figure 7B). However, there are minor differences between the two with respect to the strength of preferences for leucine, valine, glycine, and arginine. In high CAIS species, transmembrane proteins also avoid phenylalanine more strongly than non-transmembrane domains do, consistent with its role in increasing both membrane permeability and heteromeric interactions in transmembrane domains (Kwon et al., 2016; Perkins & Vaida, 2017).

**Figure 7:**
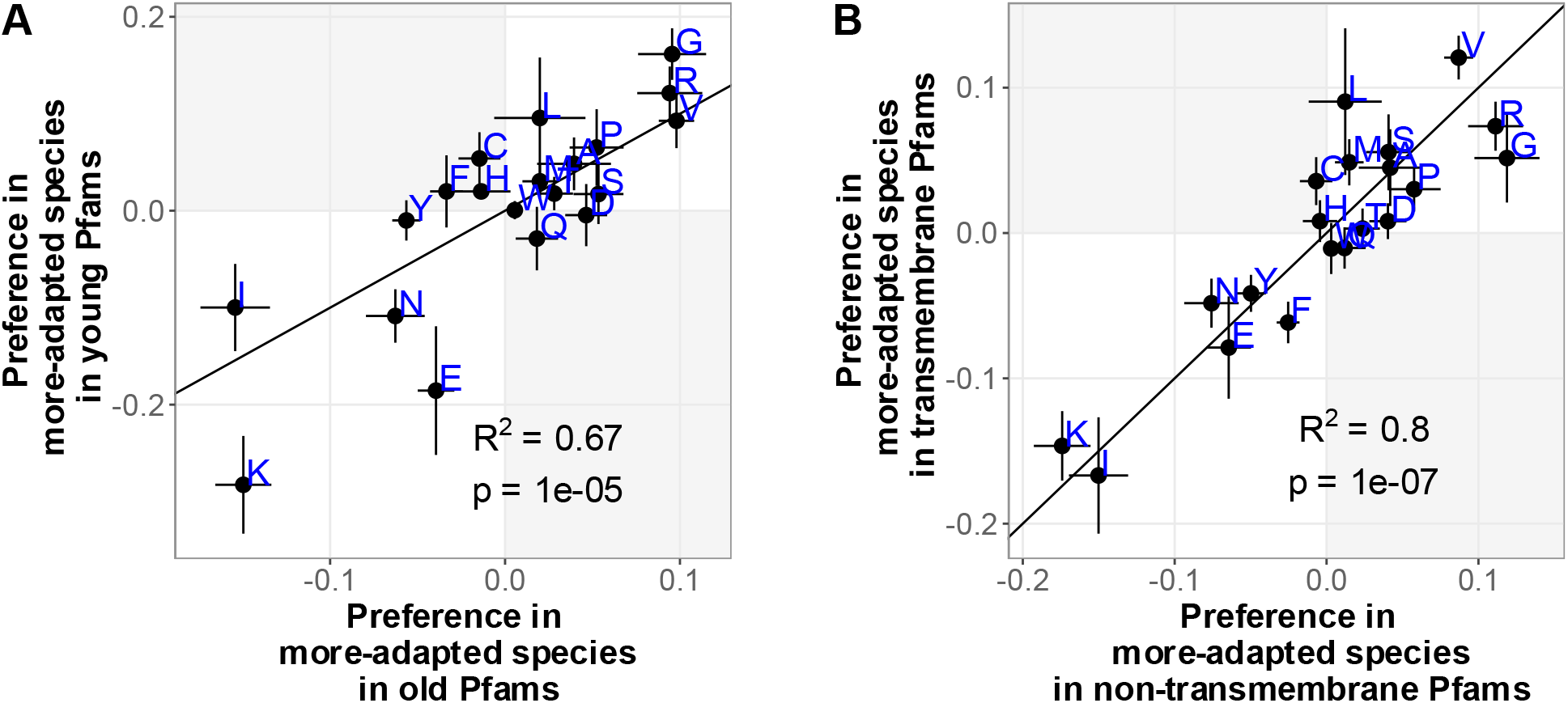
Vertebrate amino acid preferences (from our CAIS method) are similar in different groups of protein domains. Error bars show ± 1 standard error; R^2^ and p-values on figure panels are for Pearson’s correlation. Domain ages were taken from James et al. (2021), in which Pfam domains were assigned a date of evolutionary origin using TimeTree (Kumar et al., 2017). (A) Vertebrate protein domains that emerged prior to the last eukaryotic common ancestor (LECA) are categorized as “old”, and protein domains that emerged after the divergence of animals and fungi from plants as “young”. (B) Transmembrane vs non-transmembrane status was determined by James et al. (2021) using the program tmhmm (Krogh et al., 2001). In both panels, the black y=x lines indicate equal effects in the two sets of Pfam domains. Presence in an off-diagonal, shaded quadrant indicates that an amino acid trend has different signs for the two subsets of the data. Error bars indicate +/- one standard error.

### Marked preferences between evolutionarily exchangeable pairs of amino acids

Selection on subtle differences between similar amino acids may be informative regarding the biophysics of what is preferred. We selected amino acid pairs that have exchangeability values ≥ 2 in the long-timescale BLOSUM62 matrix, ≥ 1 in the short-timescale BLOSUM90 matrix (Henikoff & Henikoff, 1992), and ≥ 1 in the short-timescale PAM60 matrix (Dayhoff, 1972). These pairs are [R,K], [V,I], [D,E], [Q,E], [Y,F] and [M,L]. We further focus on the three pairs for which our three methods agree on the direction of preference: [R,K], [D,E], and [V,I]. These pairs are connected by grey lines in Figure 6. In all three cases, the CAIS method amplifies the difference within the pair to reveal a substantial preference, while the substitution model shows a difference of smaller magnitude. In all three cases (structures shown to the left of Figure 8), the side chain of the preferred amino acid has fewer degrees of rotational freedom, a biophysical difference known to impact protein stability (Cupo et al., 1980; Tang & and Dill, 1998).

**Figure 8:**
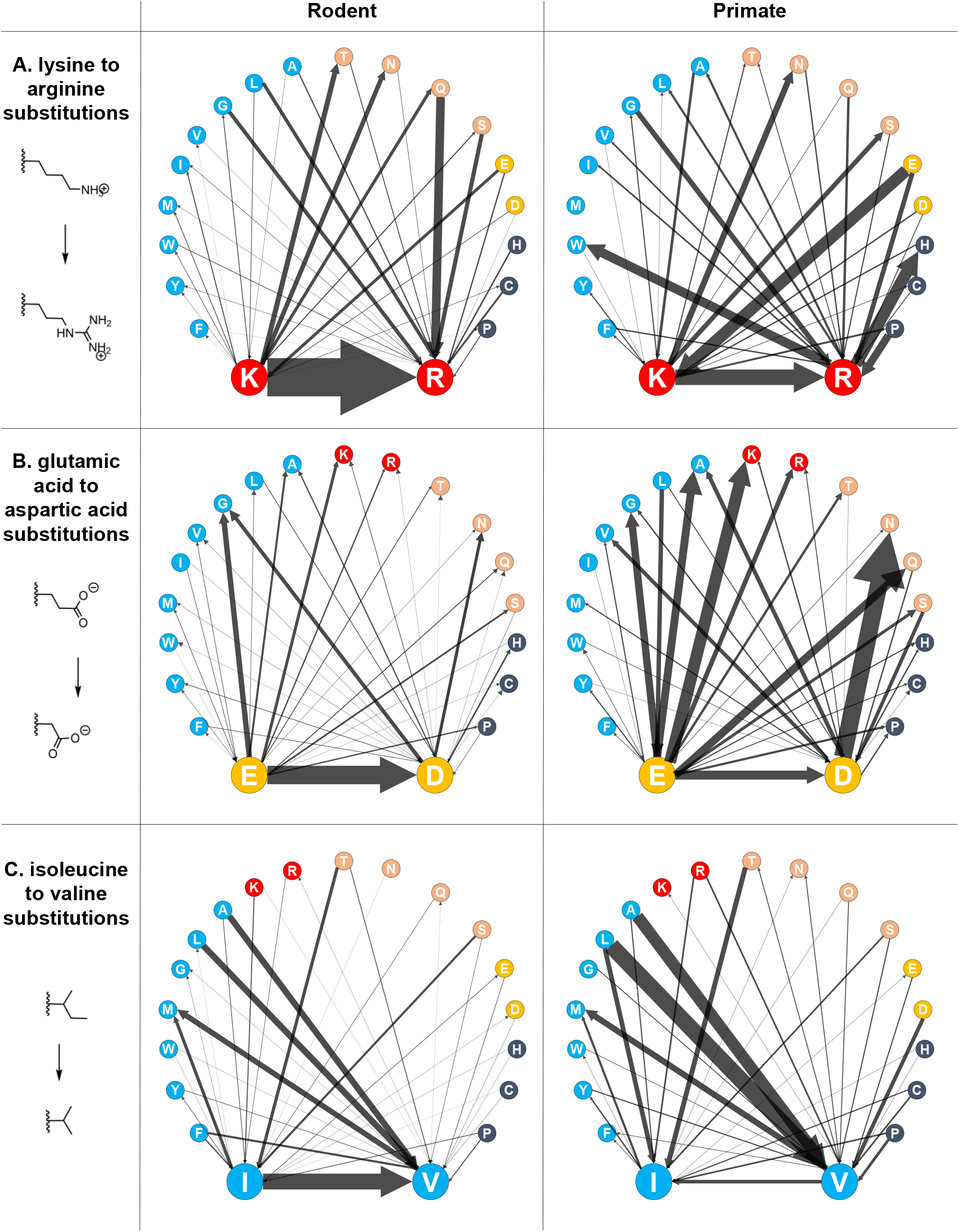
Net substitutions involving biophysically similar pairs of amino acids in rodent and primate lineages. Arrows represent all net substitutions involving (A) K and R, (B) E and D, or (C) I and V. Edge weights represent net flux between each pair (i.e. the number of K→R substitutions minus the number of R→K substitutions), normalized according to the total number of substitutions in that species pair (52,431 for rodents and 9,313 for primates). To give a sense of scale, in rodents there is 1 net substitution from K to V, 179 from Q to R, and 793 from K to R. In primates there is 1 net V→K substitution, 9 Q→R, and 57 K→R. Gross substitutions are given in Supplementary Table 4. Colors are for visual aid only and represent amino acid chemical properties: hydrophobic amino acids are blue, positively charged are red, polar non-charged are pink, negatively charged are gold, and other, “special case,” amino acids are charcoal. Note that the time non-reversibility shown should not be taken as a representation of the non-stationary amino acid frequencies – departures from stationarity are represented in Figure 6.

In the Figure 8 clamshell diagrams, we use the parsimony method to display net flux within these pairs, and from each member of the pair to the other 18 amino acids. Considering first the preference of arginine (R) over lysine (K), we note that these are the only two amino acids with sidechains that are positively charged at neutral pH. Both amino acids are basic and aliphatic, with positively charged functional groups on an alkyl side chain, and both form salt bridges and other non-covalent interactions in protein tertiary structure. The parsimony flux data shows not only that rodents gain R and lose K, but more specifically that K residues are replaced with R (Figure 8A left clamshell). Out of all ancestral rodent K residues that we observed substitutions away from, 48% became R (1452 of 3037 ancestral Ks), and 35% of derived rodent R residues came from K (1452 of 4126 derived Rs). This pattern is also found to a lesser degree in primates (113 of 526 and 705, or 21% and 16%, respectively; Figure 8A, right clamshell). Data can be found in Supplementary Table 4.

Chemical differences between the two otherwise similar amino acids may explain the preference — K has the largest median hydrophobic water-accessible surface area in folded proteins, much larger than R (47 vs. 20 Å^2^; Lins et al., 2003). In other words, at sites where an exposed K residue is replaced by an R, the exposed hydrophobic surface area is likely to be greatly reduced. This may increase the marginal stability of the folded protein. In further support of selection for stability, R can form more hydrogen bonds than K, which could have stabilizing effects enthalpically in the formation of secondary and tertiary structure, and entropically in the ordering of water molecules at the protein’s surface. Taking into account protein interactions rather than folding alone, we note that K is underrepresented in protein-protein interfaces relative to non-interface surfaces (Levy et al., 2012), while R is about equally likely to be in either. Similar considerations might also apply to protein-lipid and protein-nucleic acid interactions. Interface effects may explain the paucity of X→K substitutions: introducing K can destroy an interface and thus be strongly deleterious. X→R substitutions might be far more numerous because R is tolerated equally well in multiple contexts.

Between the two amino acids that are negatively charged at neutral pH, aspartate (D) is preferred over glutamate (E), with E→D substitutions again the predominant net flux in rodents but not primates (Figure 8B). Both amino acids bear a carboxylic acid functional group on alkyl chains of different lengths, and like R and K, both form non-covalent interactions such as salt bridges in protein tertiary structure. Like K→R substitutions, E→D substitutions entail a large reduction in hydrophobic surface area (20 → 11 Å^2^). E→D substitutions also reduce the site’s contribution to the entropic penalty of protein folding (-1.46 → -0.78 kcal/mol; Doig & Sternberg, 1995).

The small hydrophobic amino acid valine (V) is preferred over isoleucine (I), with I→V substitutions again the predominant net flux in rodents but not primates (Figure 8C). Isoleucine is very harmful in random peptides in *E. coli* (Kosinski et al., 2022), where it is more than twice as deleterious as the next most harmful amino acid. V and I have similar median hydrophobic surfaces areas (5 and 4 Å^2^, respectively). However, V side chains have a smaller entropic penalty upon protein folding than I or the structurally similar alternative, L (-0.43 vs. -0.76 vs. -0.71 kcal/mol). Such a small but fundamental effect on stability, given other similarities, could explain the preference for V as an alternative to I.

We note that in both [R,K] and [V,I] pairs, the less-preferred amino acid has an A or T nucleotide whereas the more-preferred one has a G or C (although [D,E] has evenly balanced %GC). Rodents have slightly higher genomic %GC than primates (42% vs. 41%; Weibel et al., 2024), so if the corresponding shift in mutation bias is still in process of shifting amino acid frequencies, a recent increase in %GC could contribute to preferences between similar amino acids inferred from flux. However, we observed the same preferences using our CAIS approach, which is corrected for %GC.

To better understand the effects of mutation bias, we note that they should be most pronounced at CpG sites. Four of the six R codons start with a CpG. C→T deamination at the first position would yield a cysteine (C), tryptophan (W), or stop codon, and at the second position (i.e. deamination of the C on the opposite strand) would yield a histidine (H) or glutamine (Q). Weak selection prefers R over C, W, H, and Q (Figure 6). In agreement with an expected greater relative role for mutation bias over selection in low *N*_*e*_ species, R→[C,W,H,Q] substitutions make up a larger fraction of substitutions in primates than they do in rodents (Supplementary Table 5). Net fluxes R→W and R→H are also pronounced in primates (Figure 8A).

When we consider codons with CpG in the last two positions, deamination also produces S→L, P→L, T→M, and A→V. All are counter to selective preference (Figure 6). T→M, P→L, and S→L are more common in primates than in rodents (Supplementary Table 5), and while A→V is not, this is because rodents more effectively overcome this mutation pressure with proportionately more V→A mutations (0.016 in rodents vs. 0.0099 in primates), such that A→V net flux is more pronounced in primates than rodents (Figure 8C). This demonstrates that although methyl-cytosine deamination at CpG sites is a pervasive mutational force regardless of the effectiveness of selection, more effective selection is needed to prevent the fixation of less-preferred amino acids resulting from CpG mutations.

## Discussion

Cheaper amino acids are selected for under higher *N*_*e*_. Retaining amino acids that require high GC, in the face of AT-biased mutation, also requires high *N*_*e*_. There are particularly striking preferences between highly exchangeable pairs of amino acids, with high *N*_*e*_ species preferring arginine over lysine, aspartate over glutamate, and valine over isoleucine. We find similar preferences in old vs. young and in transmembrane vs. non-transmembrane protein domains.

We used three complementary methods: two capturing amino acid composition evolution between rodents vs. primates (based on maximum likelihood and on parsimony), and one capturing amino acid frequency outcomes in a broader range of vertebrate species (based on correlation with codon adaptation as a proxy for *N*_*e*_, for which our Figure 5 analysis provides additional support). Our correlation approach is vulnerable to different sites being present in different taxa, while our flux and frequency approaches avoid this by using aligned orthologs. Parsimony flux is in theory highly vulnerable to the possibility that the effects of high *N*_*e*_ of rodents and/or the low *N*_*e*_ of primates on amino acid frequencies have already reached equilibrium. The fact that results agree with our maximum likelihood frequencies suggests that this is not the case, in agreement with literature findings of recent changes in *N*_*e*_ (Deinum et al., 2015; Jing et al., 2014; Lynch et al., 2023; Rajabi-Maham et al., 2008). Agreement between the methods, with their different biases, gives us greater confidence. Our strongest results came from maximum likelihood, with the parsimony approach providing more fine-grained information about which substitutional paths drive amino acid frequency change, and the correlation approach providing validation across a broader species range.

The parsimony flux method has been much criticized. We removed polymorphisms, which are likely to contaminate substitution counts with non-fixed slightly deleterious mutations. This improvement kept the direction but shrank the magnitude of the flux patterns observed by Jordan et al. (2005). This is compatible with shared patterns being artifacts from slightly deleterious fixations and/or a phylogenetic tree that results in better ascertainment at faster evolving sites. Whatever is responsible, the parsimony method universally produces gain of rare amino acids and loss of disorder-promoting amino acids. Slightly deleterious substitutions may be enriched for those promoting order, and/or substitutions in rapidly evolving disordered regions may be subject to a higher ascertainment rate.

Despite its weaknesses, the unique granularity of the parsimony flux method revealed that within pairs of highly exchangeable amino acids, rodents exhibit a pronounced net flux from the less favored member of the pair to the more favored, in a manner not found in primates (Figure 8). It also revealed selection’s ability to maintain CpG sites better in rodents than in primates. Our maximum likelihood method would have required an excessive number of free parameters to achieve this; it instead held the exchangeability matrix constant at values estimated for mammals as a whole. Improved methods for inferring individual substitutions (Monit & Goldstein, 2018) can avoid parsimony biases, but are potentially vulnerable to a misspecified amino acid substitution model.

A traditional view of structural biology might be skeptical of the value of information that is restricted to amino acid frequencies. However, it is striking the degree to which amino acid frequencies alone, absent information about amino acid ordering, are able to predict many properties of a polypeptide, including translational efficiency, folding kinetics, and local protein structure (Buhr et al., 2016; Saunders & Deane, 2010; Zarin et al., 2017, 2021). It is encouraging that the direction of selective preference within [K, R], [D,E], and [I, V] pairs is consistent with expected selection on protein folding and stability, as supported by differences in median hydrophobic surface area, hydrogen bonding capacity, and entropic penalty of folding.

Another line of evidence comes from the average effects of substitutions across a range of deep mutational scanning (DMS) experiments. (Sun et al., 2024) summarized the distribution of experimental fitness effects of a given substitution type with a single parameter, with small λ values being more benign. In agreement with our findings, *λ*_K→R_ = −2.77 < *λ*_R→K_ = −1.64, *λ*_E→D_ = −1.7 < *λ*_D→E_ = −1.39, and *λ*_I→V_ = −2.92 < *λ*_V→I_ = −2.64. Interestingly, the deleteriousness scores of (Alpay et al., 2025), who also analyze DMS data, show preferences for R over K and D over E. Further, they see strong preferences for V over I and D over E when inferring amino acid exchangeabilities from stability measurements alone, rather than when aggregating measurements of activity, binding, stability, expression, and fitness (see their Supplementary Figure 6).

Note that the preferences for R over K, E over D, and V over I are statistical tendencies; the opposite might still hold at (a minority of) individual sites. Because most proteins must fold (and unfold, and fold again), it is not surprising that the thermodynamics of folding would be subject to selection detectable at the level of amino acid frequencies. The specific biophysical hypotheses suggested by our observational work could be experimentally tested by aggregating data on arginine vs. lysine, aspartate over glutamate, or valine vs. isoleucine mutations across a set of model proteins, where the effect of each mutation on the thermodynamics and kinetics of folding could be measured using multiplexed assays (Atsavapranee et al., 2021; Markin et al., 2021; Tsuboyama et al., 2023).

Our observation of among-species variation in proteome-wide amino acid composition shows different patterns from previous work on within-proteome variation. Some trends are similar: like high-*N*_*e*_ species, highly expressed, slowly evolving proteins are enriched in amino acids that are cheaper to synthesize (Akashi & Gojobori, 2002). However, more generally, amino acids preferred in highly expressed proteins are not the same ones preferred under more effective selection at the species level. Highly expressed proteins are enriched in A, G, V, R, and K (Jansen & Gerstein, 2000), while slowly evolving proteins are enriched in A, G, V, D, I, M, N, Y and depleted in R (Cherry, 2010), neither of which correlates with amino acid preference under more effective selection. Disagreement may be because highly expressed proteins have different structural and functional requirements than the proteome as a whole.

Amino acid substitutions during the evolution of thermophily do, however, parallel some of our observations for more effective selection. This is expected because both more effective selection and hotter temperature are expected to favor greater protein stability. We note that proteins tend to evolve only marginal stability at mutation-selection-drift balance (Bloom, Raval, et al., 2007), with greater stability improving robustness to mistranscription and mistranslation (Drummond et al., 2005; Drummond & Wilke, 2008, 2009; Serohijos et al., 2012; Wilke et al., 2005), and potentially also mutation (Bloom, Lu, et al., 2007). In agreement with congruence between high temperature and high effective population size, Lecocq et al. (2021) found that thermophiles in the archaeal order Methanococcales use R more often, and that K→R substitutions accompany transitions to higher optimal growth temperature while R→K substitutions accompany transitions to lower growth temperature. In thermophilic species of filamentous fungi, Van Noort et al. (2013) similarly found an excess of K→R substitutions, but also a puzzling excess of D→E. Fontanillas et al. (2017) similarly found an excess of R→K, E→D, and V→I substitutions during a shift to cold adaptation of hydrothermal vent-dwelling polychaete worms. Results are thus congruent for [R,K] and [V,I], but not for [D,E]. Cold-adapted branches also had higher substitution rates and slightly higher dN/dS ratios, suggesting relaxed selection/reduced constraint relative to hot-adapted species.

Pinney et al. (2021) identified protein sites at which amino acid identity correlated with optimal growth temperature across the bacterial domain. Opposite to our work and the studies reviewed above, at these sites, K and I are strongly associated with high growth temperature in bacterial enzymes, while R and V show no association with temperature. Interestingly, hot-adapted enzymes are also enriched in I-I, K-K, and I-K spatial interactions, and K-E salt bridges are also significantly enriched in hot-adapted enzymes, while R-D salt bridges are enriched in cold-adapted enzymes (K-D and R-E salt bridges were not significant). Perhaps against a background that is generally depleted in K and I by adaptation to hot conditions, the importance of the remaining K and I interactions becomes more important. In other words, the few K and I residues remaining may have important site-specific functions.

Proteome-wide amino acid frequencies are shaped by some combination of protein domain birth, death, and tinkering. Differential rates of domain loss and duplication (James et al., 2023) shift animal proteomes toward amino acid compositions favoring lower intrinsic structural disorder (James et al., 2021). This is counterbalanced by *de novo* birth (Wilson et al., 2017). Our work here instead focuses on descent with modification, which is likely to operate on faster timescales. This focus is strictly enforced in both our parsimony and our likelihood-based flux analyses, by considering only homologous sites shared by all four species. Our outcome results with CAIS also suggest descent with modification, given that controlling for Pfam identity was not statistically supported for individual amino acids (see Methods), and that similar results were previously found for intrinsic structural disorder in which this control was included (Weibel et al., 2024).

Amino acids have distinctive biophysical properties, and selective preference among these may be universal. The concordance of our mammalian study with previous results on random peptides in *E. coli* supports the idea that for amino acid composition, “anything found to be true of *E. coli* must also be true of Elephants” (Monod & Jacob, 1961). Sometimes the demands on a position within a protein are different, e.g. for alpha helix vs. beta sheet vs. intrinsic disorder, for surface vs. buried, for acid vs. base, or for lesser vs. greater rotameric degrees of freedom, but given a particular demand, there are subtle preferences among amino acids that can deliver similar things. This is what we have quantified. This is nearly neutral theory, as applied to the essence of biochemistry.

## Materials and Methods

### Maximum likelihood equilibrium frequency estimates under a non-stationary amino acid substitution model

From the concatenated alignment of 5162 orthologous mammalian genes from (Wu et al., 2018), we retained all 16 primates, all 12 rodents, two lagomorph species that are sister to rodents (rabbit and pika) and one that is sister to primates (colugo). This results in one Glires clade (Rodents plus Lagomorphs) and one Primatomorpha clade (Primates plus colugo), as illustrated in Supplementary Figure 1.

We used the cogent3 code base (Kaehler et al., 2015; Schranz et al., 2008; Verbyla et al., 2013) to fit a non-stationary model of amino acid substitutions, given a fixed topology of the species tree taken from the maximum clade credibility consensus tree from Vertlife (Upham et al., 2019). Branch lengths were fitted by the model, but we did not model rate heterogeneity among sites. We used the QMammal exchangeability matrix (Bui et al., 2021), combined with three different equilibrium amino acid frequency vectors. One vector describes the long-term equilibrium frequency along Primatomorpha branches, another along Glires branches, and a third describes frequencies at the root. Inferred frequencies are given in Supplementary Table 1. The substitution rate matrix within a branch is then the product of the exchangeability matrix times the vector of equilibrium frequencies.

To draw out the difference between high-*N*_*e*_ Glires and low-*N*_*e*_ Primatomorpha, we first perform a logit transform on each equilibrium amino acid frequency, to reduce the heteroscedascity that is inherent to frequency measures (Warton & Hui, 2011). We then analyze differences in the logit, with (logit(Glires frequency) – logit(root frequency)) – (logit(Primatomorpha frequency) – logit(root frequency)) = Glires difference – Primatomorpha difference. We call this transformed quantity the “equilibrium frequency difference (ML)”.

### Parsimony approach to amino acid flux

We used the 65 amniote vertebrates multiple genome alignment, available on Ensembl (https://useast.ensembl.org/info/genome/compara/mlss.html?mlss=2041) (Yates et al., 2020). This alignment was generated using the Mercator-Pecan pipeline, which effectively handles rearrangements and duplications that have occurred during species’ evolutionary history by organizing genomes into segments to create a homology map, prior to alignment (Paten et al., 2008). We downloaded the aligned coding regions for our four focal species from the Ensembl REST API server (https://rest.ensembl.org/, Ensembl 98, accessed during the months of November and December 2019) using our own Python scripts, using the mouse reference genome to retrieve alignment blocks based on mouse coding regions. To be included in our dataset, genomic regions for all four of our focal species had to be present in the multiple genome alignment.

The locations of polymorphic sites to be excluded were taken from the Ensembl variation database (Hunt et al., 2018). If a polymorphism reached a threshold minor allele frequency of 5% in any of the populations in which it was reported, that site was masked in the multiple species alignment. We note that due to the larger number of studies on human populations, there are more human polymorphisms recorded in Ensembl, and thus masked in our data, and fewer for rats and chimpanzees.

Aligned DNA sequences were then translated, and informative substitutions were identified, using our own Python scripts. To be included in our analysis, an amino acid site had to be identical in three species and different in the fourth, i.e., inferred by parsimony to occur on one of the branches between mouse-rat or human-chimp. For each species in our quartet, the normalized gain/loss ratio (flux) for each amino acid was calculated by the same method as Jordan et al. (2005), where flux = (# derived – #ancestral) / (#derived + # ancestral). Net flux values and the shared and species-specific components are given in Supplementary Table 2. Data and scripts for these analyses are in the folders Parsimony_analysis and Parsimony_flux.

### Calculation of amino acid preference in vertebrates using CAIS

We use the CAIS values already calculated for 117 vertebrate species by Weibel et al. (2024). To obtain regression slopes in that paper, each species was assigned a fixed effect on the quantity of interest (in our case amino acid frequency), while controlling for Pfam identity (Mistry et al., 2021) as a random effect. While previously supported for intrinsic structural disorder, in our current analysis of amino acid frequencies, controlling for Pfam identity was not supported (p > 0.05 for all 20 amino acids). We therefore instead use the simple fractional content of each amino acid in the data for each species. We then performed 20 weighted linear regressions to predict amino acid frequencies from each species’ CAIS value, and then used the regression slopes to summarize the degree of preference for each amino acid, together with the associated standard errors on the slopes (Supplementary Table 3). Positive slopes indicate that species with more effective selection (high CAIS) prefer that amino acid, negative that they avoid it.

We do not correct for phylogeny (Felsenstein, 1985) before estimating the slopes, because phylogenetic correction amplifies differences between closely related taxa, whereas we are interested in differences that have might evolved slowly during longer periods of relatively consistent effective population size. Phylogenetic correction via phylogenetically independent contrasts (PIC) removes the relationship between CAIS slopes and other metrics.

We did this both for the whole proteome, and for the set of annotated Pfam sequences. Proteome-wide measurements yielded slopes with smaller total standard error. Where not otherwise indicated, we therefore use the proteome-wide measurements. Comparison to CAIS slopes for the subset of the proteome that belongs to a Pfam (which are subsetted in Figure 7) is shown in Supplementary Figure 4. Relevant data and scripts are in the folder CAIS.

### Pfam subsets

In the vertebrate genomes, Pfams that emerged prior to the last eukaryotic common ancestor (LECA) are identified as “old”, and Pfams that emerged after the divergence of animals and fungi from plants are identified as “young”, as annotated by James et al. (2021). Pfams were assigned as either transmembrane or non-transmembrane by James et al. (2021) using the program Tmhmm (Krogh et al., 2001). Amino acid frequencies were calculated for each Pfam subset in each species.

### Per-gene “CAIS” and CAI

We tested the relationship between CAIS and expression level, given that selection is known to be more effective for highly expressed genes (Akashi & Gojobori, 2002; Cherry, 2010). We used mouse codon frequencies from the Codon Statistics Database (K. Subramanian et al., 2022) to calculate amino acid frequencies and GC content, and used these quantities in our calculations of per-gene CAIS, as well as CAI, for all genes (ENSMUSP IDs from Ensembl 113, accessed in September 2024) 100 codons or longer for which expression data was available. Whole-organism integrated expression levels for mouse were pulled from PaxDB (Huang et al., 2023).

We calculated the CAI of a gene from relative synonymous codon usages (RSCUs) for each codon *i*, where *N*_*i*_ is the count of *i* in the gene, and *n*_*a*_ is the number of codons that code for the same amino acid as *i*:

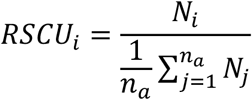

The CAI for the gene’s *L* codons, where *RSCU*_*max*_ is the maximum RSCU across all genes longer than 100 codons (1.27 for mouse, 1.35 for human), is then:

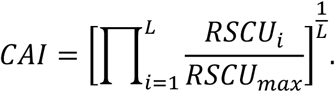

Adapting the region-specific method of Weibel et al. (2024) to calculate the “CAIS” of a single gene *g*, we first consider the expected probability *p*_*i,g*_ of seeing codon *i* in a random sequence sampled according to the observed frequency *g*_*g*_ of G or C in gene *g*, where *k*_*GC*_ is the count of G and C in the codon:

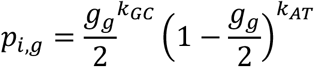

Given that amino acid *a* is found in gene *g*, the expected probability that it uses codon *i* is then:

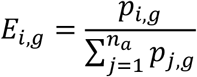

We then calculated per-gene CAIS as the Kullback-Leibler divergence of the gene’s observed codon frequencies from its expected codon probabilities:

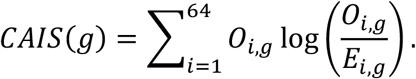

## Data availability

Full substitution tables used to calculate equilibrium frequencies, fluxes, and CAIS slopes, as well as the values themselves and all code, calculations, and supplementary data are available at https://github.com/cyclase/aa_flux. A tutorial describing the calculation of maximum likelihood amino acid frequencies under a non-stationary evolutionary model is available at https://cogent3.org/doc/examples/nonstationary_inference.html#nonstationary-model-aa-inference. All statistical modelling was done in R 3.5.1, 3.6.3, 4.2.1, or 4.3.1, and linear models were implemented specifically with the lm() function, lmodel2 (Legendre & Oksanen, 2008), or lme4 (Bates et al., 2015).

## Acknowledgments

JM, CW, and PG were supported by the John Templeton Foundation (60814 and 62220). JM, AW, PG, and GH were supported by the National Science Foundation (2333243). CW was supported by the Arnold and Mabel Beckman Foundation Scholars Program, the Western Alliance to Expand Student Opportunities (WAESO), Louis Stokes Alliance for Minority Participation (LSAMP) National Science Foundation (NSF) Cooperative Agreement (HRD-1101728), and the UA/NASA Space Grant Undergraduate Research Internship program. HM was supported by an NSF Graduate Research Fellowship. SW was supported in part by NASA Astrobiology Program ICAR grant [80NSSC21K0592] to Betül Kaçar. AW was supported by the NIH [T32 GM132008]. We thank Minh Bui, Luke Kosinski, Catherine DeBrule Stark, Maggie Horst, Sungmin Ji, and Bennett Kapili for helpful discussions. Work was partially motivated by a bet with Eugene Koonin over a fine bottle of red wine regarding the reliability of the results of Jordan et al.

## Citations

Akashi, H. (1996). Molecular Evolution Between Drosophila melanogaster and D. simulans Reduced Codon Bias, Faster Rates of Amino Acid Substitution, and Larger Proteins in D. melanogaster. Genetics, 144(3), 1297–1307. 10.1093/genetics/144.3.1297

Akashi, H., & Gojobori, T. (2002). Metabolic efficiency and amino acid composition in the proteomes of Escherichia coli and Bacillus subtilis. Proceedings of the National Academy of Sciences, 99(6), 3695–3700. 10.1073/pnas.062526999

Alpay, B. A., Nanda, P., Nagy, E., & Desai, M. M. (2025). Amino acid exchangeability and surface accessibility underpin the effects of single substitutions (p. 2025.06.13.659595). bioRxiv. 10.1101/2025.06.13.659595

Atsavapranee, B., Stark, C. D., Sunden, F., Thompson, S., & Fordyce, P. M. (2021). Fundamentals to function: Quantitative and scalable approaches for measuring protein stability. Cell Systems, 12(6), 547–560. 10.1016/j.cels.2021.05.009

Bates, D., Mächler, M., Bolker, B., & Walker, S. (2015). Fitting Linear Mixed-Effects Models Using lme4. Journal of Statistical Software, 67, 1–48. 10.18637/jss.v067.i01

Bénitière, F., Lefébure, T., & Duret, L. (2025). Variation in the fitness impact of translationally optimal codons among animals. Genome Research, gr.279837.124. 10.1101/gr.279837.124

Bloom, J. D., Lu, Z., Chen, D., Raval, A., Venturelli, O. S., & Arnold, F. H. (2007). Evolution favors protein mutational robustness in sufficiently large populations. BMC Biology, 5(1), 29. 10.1186/1741-7007-5-29

Bloom, J. D., Raval, A., & Wilke, C. O. (2007). Thermodynamics of neutral protein evolution. Genetics, 175(1), 255–266. 10.1534/genetics.106.061754

Buhr, F., Jha, S., Thommen, M., Mittelstaet, J., Kutz, F., Schwalbe, H., Rodnina, M. V., & Komar, A. A. (2016). Synonymous Codons Direct Cotranslational Folding toward Different Protein Conformations. Molecular Cell, 61(3), 341–351. 10.1016/j.molcel.2016.01.008

Bui, M. Q., Dang, C. C., Vinh, L. S., & Lanfear, R. (2021). QMaker: Fast and Accurate Method to Estimate Empirical Models of Protein Evolution. Systematic Biology, 70(5), 1046–1060. 10.1093/sysbio/syab010

Cherry, J. L. (2010). Highly expressed and slowly evolving proteins share compositional properties with thermophilic proteins. Molecular Biology and Evolution, 27(3), 735–741. 10.1093/molbev/msp270

Costa, I. R., Thompson, J. D., Ortega, J. M., & Prosdocimi, F. (2015). Metazoan Remaining Genes for Essential Amino Acid Biosynthesis: Sequence Conservation and Evolutionary Analyses. Nutrients, 7(1), Article 1. 10.3390/nu7010001

Cupo, P., El-Deiry, W., Whitney, P. L., & Awad, W. M. (1980). Stabilization of proteins by guanidination. Journal of Biological Chemistry, 255(22), 10828–10833. 10.1016/S0021-9258(19)70382-2

Dang, C. C., Minh, B. Q., McShea, H., Masel, J., James, J. E., Vinh, L. S., & Lanfear, R. (2022). nQMaker: Estimating Time Nonreversible Amino Acid Substitution Models. Systematic Biology, syac007. 10.1093/sysbio/syac007

Dayhoff, M. O. (1972). Atlas of Protein Sequence and Structure. National Biomedical Research Foundation.

Deinum, E. E., Halligan, D. L., Ness, R. W., Zhang, Y.-H., Cong, L., Zhang, J.-X., & Keightley, P. D. (2015). Recent Evolution in Rattus norvegicus Is Shaped by Declining Effective Population Size. Molecular Biology and Evolution, 32(10), 2547–2558. 10.1093/molbev/msv126

Doig, A. J., & Sternberg, M. J. E. (1995). Side-chain conformational entropy in protein folding. Protein Science, 4(11), 2247–2251. 10.1002/pro.5560041101

Drummond, D. A., Bloom, J. D., Adami, C., Wilke, C. O., & Arnold, F. H. (2005). Why highly expressed proteins evolve slowly. Proceedings of the National Academy of Sciences, 102(40), 14338–14343. 10.1073/pnas.0504070102

Drummond, D. A., & Wilke, C. O. (2008). Mistranslation-Induced Protein Misfolding as a Dominant Constraint on Coding-Sequence Evolution. Cell, 134(2), 341–352. 10.1016/j.cell.2008.05.042

Drummond, D. A., & Wilke, C. O. (2009). The evolutionary consequences of erroneous protein synthesis. Nature Reviews Genetics, 10(10), Article 10. 10.1038/nrg2662

Eőry, L., Hllign, D.L., & Keightley, P. D. (2010). Distributions of Selectively Constr ined Sites nd Deleterious Mutation Rates in the Hominid and Murid Genomes. Molecular Biology and Evolution, 27(1), 177–192. 10.1093/molbev/msp219

Felsenstein, J. (1985). Phylogenies and the Comparative Method. The American Naturalist, 125(1), 1–15. 10.1086/284325

Fontanillas, E., Galzitskaya, O. V., Lecompte, O., Lobanov, M. Y., Tanguy, A., Mary, J., Girguis, P. R., Hourdez, S., & Jollivet, D. (2017). Proteome Evolution of Deep-Sea Hydrothermal Vent Alvinellid Polychaetes Supports the Ancestry of Thermophily and Subsequent Adaptation to Cold in Some Lineages. Genome Biology and Evolution, 9(2), 279–296. 10.1093/gbe/evw298

Galtier, N., Roux, C., Rousselle, M., Romiguier, J., Figuet, E., Glémin, S., Bierne, N., & Duret, L. (2018). Codon Usage Bias in Animals: Disentangling the Effects of Natural Selection, Effective Population Size, and GC-Biased Gene Conversion. Molecular Biology and Evolution, 35(5), 1092–1103. 10.1093/molbev/msy015

Goldstein, R. A., & Pollock, D. D. (2006). Observations of Amino Acid Gain and Loss during Protein Evolution Are Explained by Statistical Bias. Molecular Biology and Evolution, 23(7), 1444–1449. 10.1093/molbev/msl010

Groussin, M., Boussau, B., & Gouy, M. (2013). A branch-heterogeneous model of protein evolution for efficient inference of ancestral sequences. Systematic Biology, 62(4), 523–538. 10.1093/sysbio/syt016

Haag-Liautard, C., Coffey, N., Houle, D., Lynch, M., Charlesworth, B., & Keightley, P. D. (2008). Direct Estimation of the Mitochondrial DNA Mutation Rate in Drosophila melanogaster. PLoS Biology, 6(8), e204. 10.1371/journal.pbio.0060204

Halligan, D. L., Oliver, F., Eyre-Walker, A., Harr, B., & Keightley, P. D. (2010). Evidence for Pervasive Adaptive Protein Evolution in Wild Mice. PLOS Genetics, 6(1), e1000825. 10.1371/journal.pgen.1000825

Henikoff, S., & Henikoff, J. G. (1992). Amino acid substitution matrices from protein blocks. Proceedings of the National Academy of Sciences of the United States of America, 89(22), 10915–10919. 10.1073/pnas.89.22.10915

Hershberg, R., & Petrov, D. A. (2010). Evidence That Mutation Is Universally Biased towards AT in Bacteria. PLoS Genetics, 6(9), e1001115. 10.1371/journal.pgen.1001115

Huang, Q., Szklarczyk, D., Wang, M., Simonovic, M., & von Mering, C. (2023). PaxDb 5.0: Curated Protein Quantification Data Suggests Adaptive Proteome Changes in Yeasts. Molecular & Cellular Proteomics, 22(10), 100640. 10.1016/j.mcpro.2023.100640

Hunt, S. E., McLaren, W., Gil, L., Thormann, A., Schuilenburg, H., Sheppard, D., Parton, A., Armean, I. M., Trevanion, S. J., Flicek, P., & Cunningham, F. (2018). Ensembl variation resources. Database, 2018, bay119. 10.1093/database/bay119

Hurst, L. D., Feil, E. J., & Rocha, E. P. C. (2006). Causes of trends in amino-acid gain and loss. Nature, 442(7105), E11–E12. 10.1038/nature05137

James, J. E., Nelson, P., & Masel, J. (2023). Differential retention of Pfam domains creates long-term evolutionary trends. Molecular Biology and Evolution, 2023. 10.1101/2022.10.27.514087

James, J. E., Willis, S. M., Nelson, P. G., Weibel, C., Kosinski, L. J., & Masel, J. (2021). Universal and taxon-specific trends in protein sequences as a function of age. eLife, 10, e57347. 10.7554/eLife.57347

Jansen, R., & Gerstein, M. (2000). Analysis of the yeast transcriptome with structural and functional categories: Characterizing highly expressed proteins. Nucleic Acids Research, 28(6), 1481–1488. 10.1093/nar/28.6.1481

Jing, M., Yu, H.-T., Bi, X., Lai, Y.-C., Jiang, W., & Huang, L. (2014). Phylogeography of Chinese house mice (Mus musculus musculus/castaneus): Distribution, routes of colonization and geographic regions of hybridization. Molecular Ecology, 23(17), 4387–4405. 10.1111/mec.12873

Jordan, I. K., Kondrashov, F. A., Adzhubei, I. A., Wolf, Y. I., Koonin, E. V., Kondrashov, A. S., & Sunyaev, S. (2005). A universal trend of amino acid gain and loss in protein evolution. Nature, 433(7026), 633–638. 10.1038/nature03306

Kaehler, B. D., Yap, V. B., Zhang, R., & Huttley, G. A. (2015). Genetic Distance for a General Non-Stationary Markov Substitution Process. Systematic Biology, 64(2), 281–293. 10.1093/sysbio/syu106

Keightley, P. D., Lercher, M. J., & Eyre-Walker, A. (2005). Evidence for Widespread Degradation of Gene Control Regions in Hominid Genomes. PLOS Biology, 3(2), e42. 10.1371/journal.pbio.0030042

Kosinski, L., Aviles, N., Gomez, K., & Masel, J. (2022). Random peptides rich in small and disorder-promoting amino acids are less likely to be harmful. Genome Biology and Evolution, evac085. 10.1093/gbe/evac085

Krogh, A., Larsson, B., von Heijne, G., & Sonnhammer, E. L. L. (2001). Predicting transmembrane protein topology with a hidden markov model: Application to complete genomes. Journal of Molecular Biology, 305(3), 567–580. 10.1006/jmbi.2000.4315

Kumar, S., Stecher, G., Suleski, M., & Hedges, S. B. (2017). TimeTree: A Resource for Timelines, Timetrees, and Divergence Times. Molecular Biology and Evolution, 34(7), 1812–1819. 10.1093/molbev/msx116

Kwon, M. J., Park, J., Jang, S., Eom, C. Y., & Oh, E. S. (2016). The conserved phenylalanine in the transmembrane domain enhances heteromeric interactions of syndecans. Journal of Biological Chemistry, 291(2), 872–881. 10.1074/jbc.M115.685040

Le, S. Q., Dang, C. C., & Gascuel, O. (2012). Modeling Protein Evolution with Several Amino Acid Replacement Matrices Depending on Site Rates. Molecular Biology and Evolution, 29(10), 2921– 2936. 10.1093/molbev/mss112

Le, S. Q., & Gascuel, O. (2008). An improved general amino acid replacement matrix. Molecular Biology and Evolution, 25(7), 1307–1320. 10.1093/molbev/msn067

Lecocq, M., Groussin, M., Gouy, M., & Brochier-Armanet, C. (2021). The Molecular Determinants of Thermoadaptation: Methanococcales as a Case Study. Molecular Biology and Evolution, 38(5), 1761–1776. 10.1093/molbev/msaa312

Legendre, P., & Oksanen, J. (2008). lmodel2: Model II Regression (p. 1.7-4) [Computer software]. https://CRAN.R-project.org/package=lmodel2

Levy, E. D., De, S., & Teichmann, S. A. (2012). Cellular crowding imposes global constraints on the chemistry and evolution of proteomes. Proceedings of the National Academy of Sciences, 109(50), 20461–20466. 10.1073/pnas.1209312109

Lins, L., Thomas, A., & Brasseur, R. (2003). Analysis of accessible surface of residues in proteins. Protein Science, 12(7), 1406–1417. 10.1110/ps.0304803

Lynch, M. (2010). Rate, molecular spectrum, and consequences of human mutation. Proceedings of the National Academy of Sciences, 107(3), 961–968. 10.1073/pnas.0912629107

Lynch, M., Ali, F., Lin, T., Wang, Y., Ni, J., & Long, H. (2023). The divergence of mutation rates and spectra across the Tree of Life. EMBO Reports, 24(10), e57561. 10.15252/embr.202357561

Lynch, M., & Conery, J. S. (2003). The Origins of Genome Complexity. Science, 302(5649), 1401–1404. 10.1126/science.1089370

Lynch, M., Sung, W., Morris, K., Coffey, N., Landry, C. R., Dopman, E. B., Dickinson, W. J., Okamoto, K., Kulkarni, S., Hartl, D. L., & Thomas, W. K. (2008). A genome-wide view of the spectrum of spontaneous mutations in yeast. Proceedings of the National Academy of Sciences of the United States of America, 105(27), 9272. 10.1073/pnas.0803466105

Markin, C. J., Mokhtari, D. A., Sunden, F., Appel, M. J., Akiva, E., Longwell, S. A., Sabatti, C., Herschlag, D., & Fordyce, P. M. (2021). Revealing enzyme functional architecture via high-throughput microfluidic enzyme kinetics. Science, 373(6553), eabf8761. 10.1126/science.abf8761

McDonald, J. H. (2006). Apparent Trends of Amino Acid Gain and Loss in Protein Evolution Due to Nearly Neutral Variation. Molecular Biology and Evolution, 23(2), 240–244. 10.1093/molbev/msj026

Mistry, J., Chuguransky, S., Williams, L., Qureshi, M., Salazar, G. A., Sonnhammer, E. L. L., Tosatto, S. C. E., Paladin, L., Raj, S., Richardson, L. J., Finn, R. D., & Bateman, A. (2021). Pfam: The protein families database in 2021. Nucleic Acids Research, 49(D1), D412–D419. 10.1093/nar/gkaa913

Monit, C., & Goldstein, R. A. (2018). SubRecon: Ancestral reconstruction of amino acid substitutions along a branch in a phylogeny. Bioinformatics, 34(13), 2297–2299. 10.1093/bioinformatics/bty101

Monod, J., & Jacob, F. (1961). Teleonomic mechanisms in cellular metabolism, growth, and differentiation. Cold Spring Harbor Symposia on Quantitative Biology, 26, 389–401. 10.1101/sqb.1961.026.01.048

Muñoz-Gómez, S. A., Susko, E., Williamson, K., Eme, L., Slamovits, C. H., Moreira, D., López-García, P., & Roger, A. J. (2022). Site-and-branch-heterogeneous analyses of an expanded dataset favour mitochondria as sister to known Alphaproteobacteria. Nature Ecology & Evolution, 6(3), 253– 262. 10.1038/s41559-021-01638-2

Neme, R., Amador, C., Yildirim, B., McConnell, E., & Tautz, D. (2017). Random sequences are an abundant source of bioactive RNAs or peptides. Nature Ecology and Evolution, 1(6). 10.1038/s41559-017-0127

Ohta, T. (1973). Slightly Deleterious Mutant Substitutions in Evolution. Nature, 246(5428), 96–98. 10.1038/246096a0

Paten, B., Herrero, J., Beal, K., Fitzgerald, S., & Birney, E. (2008). Enredo and Pecan: Genome-wide mammalian consistency-based multiple alignment with paralogs. Genome Research, 18(11), 1814–1828. 10.1101/gr.076554.108

Perkins, R., & Vaida, V. (2017). Phenylalanine Increases Membrane Permeability. Journal of the American Chemical Society, 139(41), 14388–14391. 10.1021/jacs.7b09219

Phifer-Rixey, M., Bonhomme, F., Boursot, P., Churchill, G. A., Piálek, J., Tucker, P. K., & Nachman, M. W. (2012). Adaptive Evolution and Effective Population Size in Wild House Mice. Molecular Biology and Evolution, 29(10), 2949–2955. 10.1093/molbev/mss105

Pinney, M. M., Mokhtari, D. A., Akiva, E., Yabukarski, F., Sanchez, D. M., Liang, R., Doukov, T., Martinez, T. J., Babbitt, P. C., & Herschlag, D. (2021). Parallel molecular mechanisms for enzyme temperature adaptation. Science, 371(6533), eaay2784. 10.1126/science.aay2784

Raiford, D. W., Heizer, E. M., Miller, R. V., Akashi, H., Raymer, M. L., & Krane, D. E. (2008). Do Amino Acid Biosynthetic Costs Constrain Protein Evolution in Saccharomyces cerevisiae? Journal of Molecular Evolution, 67(6), 621–630. 10.1007/s00239-008-9162-9

Rajabi-Maham, H., Orth, A., & Bonhomme, F. (2008). Phylogeography and postglacial expansion of Mus musculus domesticus inferred from mitochondrial DNA coalescent, from Iran to Europe. Molecular Ecology, 17(2), 627–641. 10.1111/j.1365-294X.2007.03601.x

Saunders, R., & Deane, C. M. (2010). Synonymous codon usage influences the local protein structure observed. Nucleic Acids Research, 38(19), 6719–6728. 10.1093/nar/gkq495

Schranz, H. W., Yap, V. B., Easteal, S., Knight, R., & Huttley, G. A. (2008). Pathological rate matrices: From primates to pathogens. BMC Bioinformatics, 9(1), 550. 10.1186/1471-2105-9-550

Serohijos, A. W. R., Rimas, Z., & Shakhnovich, E. I. (2012). Protein Biophysics Explains Why Highly Abundant Proteins Evolve Slowly. Cell Reports, 2(2), 249–256. 10.1016/j.celrep.2012.06.022

Sharp, P. M., Bailes, E., Grocock, R. J., Peden, J. F., & Sockett, R. E. (2005). Variation in the strength of selected codon usage bias among bacteria. Nucleic Acids Research, 33(4), 1141–1153. 10.1093/nar/gki242

Sharp, P. M., Emery, L. R., & Zeng, K. (2010). Forces that influence the evolution of codon bias. Philosophical Transactions of the Royal Society B: Biological Sciences, 365(1544), 1203–1212. 10.1098/rstb.2009.0305

Sharp, P. M., & Li, W.-H. (1987). The codon adaptation index-a measure of directional synonymous codon usage bias, and its potential applications. Nucleic Acids Research, 15(3), 1281–1295. 10.1093/nar/15.3.1281

Subramanian, K., Payne, B., Feyertag, F., & Alvarez-Ponce, D. (2022). The Codon Statistics Database: A Database of Codon Usage Bias. Molecular Biology and Evolution, 39(8), msac157. 10.1093/molbev/msac157

Subramanian, S. (2008). Nearly neutrality and the evolution of codon usage bias in eukaryotic genomes. Genetics, 178(4), 2429–2432. 10.1534/genetics.107.086405

Sun, M., Stoltzfus, A., & McCandlish, D. M. (2024). A fitness distribution law for amino-acid replacements (p. 2024.10.11.617952). bioRxiv. 10.1101/2024.10.11.617952

Tang, K. E. S., & and Dill, K. A. (1998). Native Protein Fluctuations: The Conformational-Motion Temperature and the Inverse Correlation of Protein Flexibility with Protein Stability. Journal of Biomolecular Structure and Dynamics, 16(2), 397–411. 10.1080/07391102.1998.10508256

Tenesa, A., Navarro, P., Hayes, B. J., Duffy, D. L., Clarke, G. M., Goddard, M. E., & Visscher, P. M. (2007). Recent human effective population size estimated from linkage disequilibrium. Genome Research, 17(4), 520–526. 10.1101/gr.6023607

Theillet, F.-X., Kalmar, L., Tompa, P., Han, K.-H., Selenko, P., Dunker, A. K., Daughdrill, G. W., & Uversky, V. N. (2013). The alphabet of intrinsic disorder. Intrinsically Disordered Proteins, 1(1), e24360. 10.4161/idp.24360

Tien, M. Z., Meyer, A. G., Sydykova, D. K., Spielman, S. J., & Wilke, C. O. (2013). Maximum Allowed Solvent Accessibilites of Residues in Proteins. PLOS ONE, 8(11), e80635. 10.1371/journal.pone.0080635

Trifonov, E. N. (2000). Consensus temporal order of amino acids and evolution of the triplet code. Gene, 261(1), 139–151. 10.1016/S0378-1119(00)00476-5

Tsuboyama, K., Dauparas, J., Chen, J., Laine, E., Mohseni Behbahani, Y., Weinstein, J. J., Mangan, N. M., Ovchinnikov, S., & Rocklin, G. J. (2023). Mega-scale experimental analysis of protein folding stability in biology and design. Nature, 620(7973), 434–444. 10.1038/s41586-023-06328-6

Upham, N. S., Esselstyn, J. A., & Jetz, W. (2019). Inferring the mammal tree: Species-level sets of phylogenies for questions in ecology, evolution, and conservation. PLOS Biology, 17(12), e3000494. 10.1371/journal.pbio.3000494

van Noort, V., Bradatsch, B., Arumugam, M., Amlacher, S., Bange, G., Creevey, C., Falk, S., Mende, D. R., Sinning, I., Hurt, E., & Bork, P. (2013). Consistent mutational paths predict eukaryotic thermostability. BMC Evolutionary Biology, 13(1), 7. 10.1186/1471-2148-13-7

Van Oss, S. B., & Carvunis, A.-R. (2019). De novo gene birth. PLOS Genetics, 15(5), e1008160. 10.1371/journal.pgen.1008160

Verbyla, K. L., Yap, V. B., Pahwa, A., Shao, Y., & Huttley, G. A. (2013). The Embedding Problem for Markov Models of Nucleotide Substitution. PLOS ONE, 8(7), e69187. 10.1371/journal.pone.0069187

Warton, D. I., & Hui, F. K. C. (2011). The arcsine is asinine: The analysis of proportions in ecology. Ecology, 92(1), 3–10. 10.1890/10-0340.1

Wehbi, S., Wheeler, A., Morel, B., Manepalli, N., Minh, B. Q., Lauretta, D. S., & Masel, J. (2024). Order of mino cid recruitment into the genetic code resolved by l st univers l common ncestor’s protein domains. Proceedings of the National Academy of Sciences, 121(52), e2410311121. 10.1073/pnas.2410311121

Weibel, C. A., Wheeler, A. L., James, J. E., Willis, S. M., McShea, H., & Masel, J. (2024). The protein domains of vertebrate species in which selection is more effective have greater intrinsic structural disorder. eLife, 12, RP87335. 10.7554/eLife.87335

Whelan, S., & Goldman, N. (2001). A General Empirical Model of Protein Evolution Derived from Multiple Protein Families Using a Maximum-Likelihood Approach. Molecular Biology and Evolution, 18(5), 691–699. 10.1093/oxfordjournals.molbev.a003851

Wilke, C. O., Bloom, J. D., Drummond, D. A., & Raval, A. (2005). Predicting the Tolerance of Proteins to Random Amino Acid Substitution. Biophysical Journal, 89(6), 3714–3720. 10.1529/biophysj.105.062125

Wilson, B. A., Foy, S. G., Neme, R., & Masel, J. (2017). Young genes are highly disordered as predicted by the preadaptation hypothesis of de novo gene birth. Nature Ecology & Evolution, 1(6), Article 6. 10.1038/s41559-017-0146

Wu, S., Edwards, S., & Liu, L. (2018). Genome-scale DNA sequence data and the evolutionary history of placental mammals. Data in Brief, 18, 1972–1975. 10.1016/j.dib.2018.04.094

Yang, Z., & Nielsen, R. (2008). Mutation-Selection Models of Codon Substitution and Their Use to Estimate Selective Strengths on Codon Usage. Molecular Biology and Evolution, 25(3), 568–579. 10.1093/molbev/msm284

Yates, A. D., Achuthan, P., Akanni, W., Allen, J., Allen, J., Alvarez-Jarreta, J., Amode, M. R., Armean, I. M., Azov, A. G., Bennett, R., Bhai, J., Billis, K., Boddu, S., Marugán, J. C., Cummins, C., Davidson, C., Dodiy, K., Ftim, R.G ll, A., … Flicek, P. (2020). Ensembl 2020. Nucleic Acids Research, 48(D1), D682–D688. 10.1093/nar/gkz966

Zrin, T., Strome, B., Peng, G., Pritišnc, I., Formn-Kay, J.D., & Moses, A. M. (2021). Identifying molecular features that are associated with biological function of intrinsically disordered protein regions. eLife, 10, e60220. 10.7554/eLife.60220

Zarin, T., Tsai, C. N., Nguyen Ba, A. N., & Moses, A. M. (2017). Selection maintains signaling function of a highly diverged intrinsically disordered region. Proceedings of the National Academy of Sciences, 114(8), E1450–E1459. 10.1073/pnas.1614787114

Zuckerkandl, E., Derancourt, J., & Vogel, H. (1971). Mutational trends and random processes in the evolution of informational macromolecules. Journal of Molecular Biology, 59(3), 473–490. 10.1016/0022-2836(71)90311-1

