## Supplementary Figures for "The effectiveness of selection in a species affects the direction of amino acid frequency evolution"

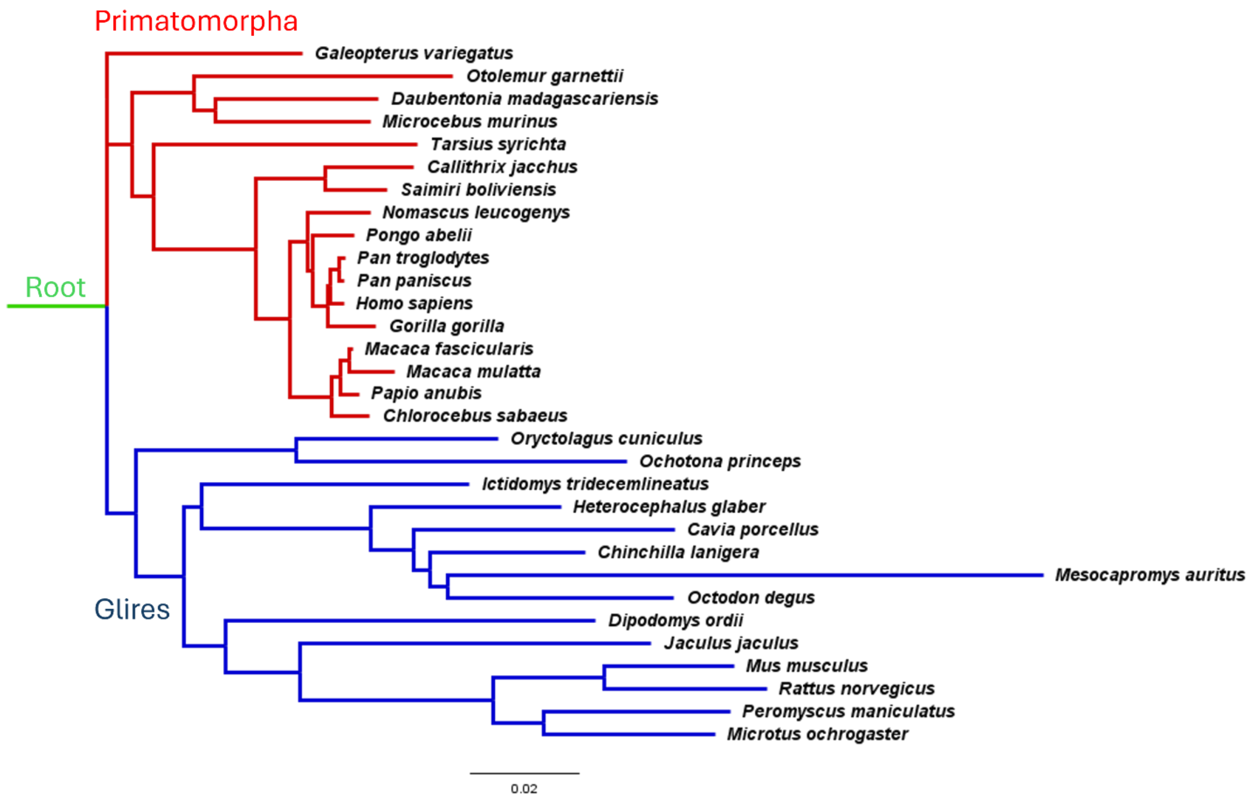


**Supplementary Figure 1:** Phylogeny used to calculate nonstationary amino acid frequency vectors. Primatomorpha includes primates and sister species Sunda flying lemur (*Galeopteris variegatus*), and Glires includes rodents as well as sister species European rabbit (*Oryctolagus cuniculus*) and American pika (*Ochotona princeps*). Branch lengths reflect expected number of substitutions per site as fitted by cogent3. Species tree topology is the maximum clade credibility consensus tree from Vertlife (Upham et al., 2019).

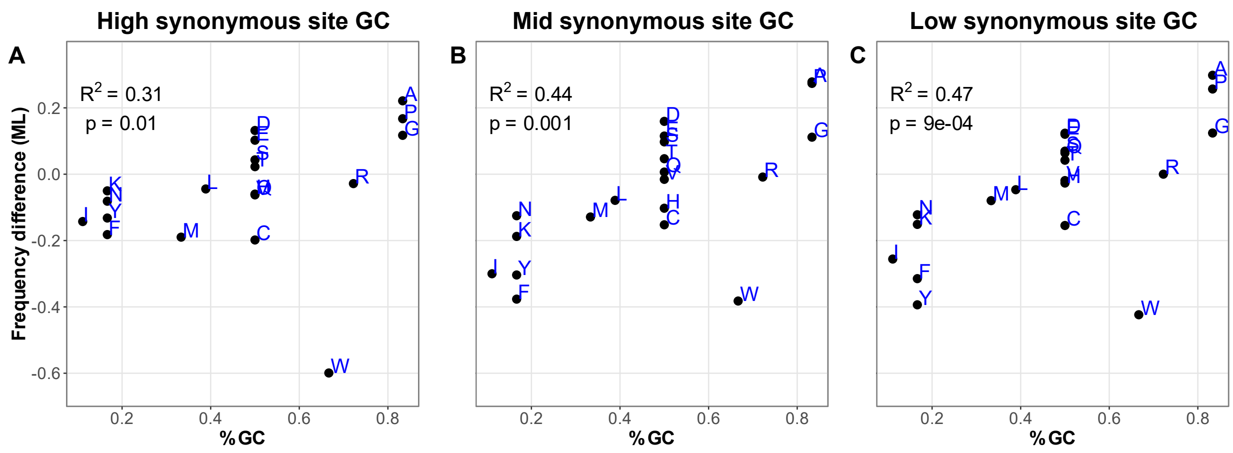


**Supplementary Figure 2:** Differences between Glires and Primatomorpha (see Figure 2B) are more pronounced when synonymous site GC content is lower. Here, we calculated a gene's synonymous site GC as the count of codons with G or C at a degenerate site over the total number of codons for that amino acid, including both first and third position degeneracy. For example, synonymous site GC for valine alone would be equal to (#GTG + #GTC) / (#GTG + #GTC + #GTA + #GTT); the full formula is (GTG+GTC+GCG+GCC+GAC+GAG+GGG+GGC+TTC+TTG+CTC+CTG+TCC+TCG+AGC+TAC+TAG+TGC+CCC+CCG+CAC+CAG+CGC+CGG+AGG+ATC+ACC+ACG+AAC+AAG+CTG+CTA+CGG+CGA)/(GTG+GTC+GTA+GTT+GCG+GCC+GCA+GCT+GAC+GAT+GAG+GAA+GGG+GGC+GGA+GGT+TTC+TTT+TTG+TTA+CTC+CTG+CTA+CTT+TCC+TCG+TCA+TCT+AGC+AGT+TAC+TAT+TAG+TAA+TGC+TGT+CCC+CCG+CCA+CCT+CAC+CAT+CAG+CAA+CGC+CGG+CGA+CGT+AGG+AGA+ATC+ATA+ATT+ACC+ACG+ACA+ACT+AAC+AAT+AAG+AAA+CTG+TTG+CTA+TTA+CGG+AGG+CGA+AGA). Genes were then split into three equally sized bins based on human synonymous GC content, and maximum likelihood equilibrium frequencies were calculated for Primatomorpha, Glires, and root from the concatenated alignment of each bin. All data and scripts for maximum likelihood analysis, can be found in the ML_flux folder of the project GitHub page. $R^2$ here uses Spearman's R, rather than Pearson's as used elsewhere in this manuscript, in order to minimize the impact of variation in the degree to which tryptophan (W) acts as an outlier. As in Figure 2, ML frequencies were logit transformed before the difference was taken.

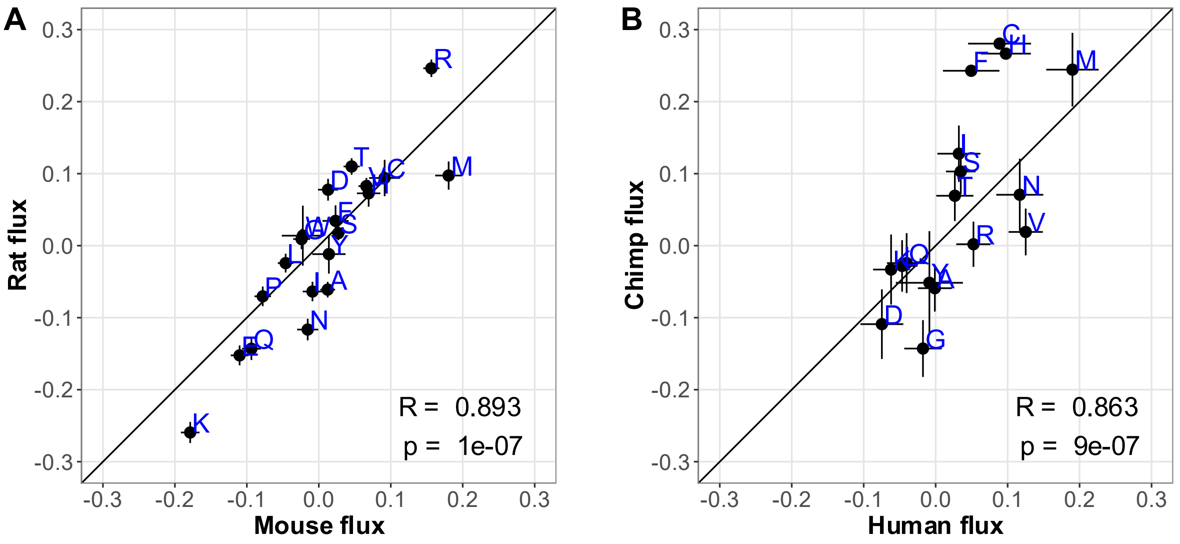


**Supplementary Figure 3**: Closely related species have similar amino acid fluxes; a tendency toward larger magnitude fluxes for rat than for mouse (A), and for chimp than for human (B), might reflect relative data quality. Diagonal lines show y = x; error bars show standard errors; R and p-values on figure panels are for Pearson's R correlation.

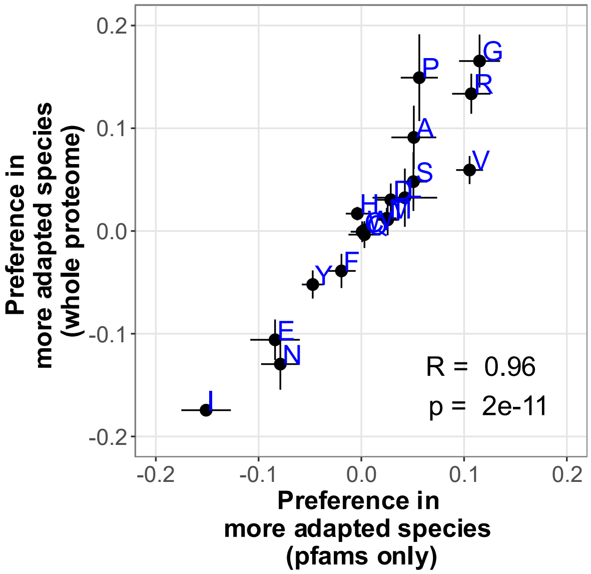


**Supplementary Figure 4**: CAIS slopes are similar whether calculated in Pfam domains or across the whole proteome. Error bars show ± 1 standard error; R and P-values on figure panels are for Pearson's R correlation.

### References

Upham, N. S., Esselstyn, J. A., & Jetz, W. (2019). Inferring the mammal tree: Species-level sets of phylogenies for questions in ecology, evolution, and conservation. *PLOS Biology*, *17*(12), e3000494. https://doi.org/10.1371/journal.pbio.3000494
